# Ancient origin of the dorso-ventral patterning system of vertebrate paired fins

**DOI:** 10.64898/2026.02.08.704681

**Authors:** Rebecca E. Dale, Silke Berger, Sara Alaei, Adele Barugahare, Marcus C. Davis, Laura Perlaza-Jimenez, Frank J. Tulenko, Peter D. Currie

## Abstract

The origin of paired fins was a major event in early vertebrate history that fuelled the adaptive radiation of the gnathostome clade. Evidence for the conservation of ancient regulatory systems between paired and median fins supports a model in which appendage-patterning mechanisms first evolved in the midline and were later redeployed to a new embryonic context, the flank. However, while paired fins and limbs are asymmetric along the dorso-ventral (DV) axis, the equivalent axis does not exist in median fins, which are symmetric across the midline, raising questions of how DV patterning evolved. Here, we combine genetic tools in zebrafish with comparative expression analyses in representative ray-finned and cartilaginous fishes to show that a subset of limb dorsoventral (DV) patterning genes are expressed during median fin development. Using genetic lineage tracing of the canonical limb dorsalizing factor *lmx1bb*, we further demonstrate that, although *lmx1bb*-derived cells occupy distinct spatial domains across fin types, a subset of this lineage is fated to differentiate into fin-ray osteoblasts in both paired and median fins. To test the potential conservation of regulatory modules across fin types, we generated a series of deletion mutants for two known *lmx1b* mouse limb enhancers, *LARM1* and *LARM2,* that we show are partially conserved upstream of zebrafish *lmx1bb*. These experiments revealed that *LARM* function is essential for dorsoventral patterning in the paired pectoral and pelvic fins of zebrafish but is dispensable for formation of the unpaired dorsal and anal fins. To determine if the paired fin specificity of *LARM*-mediated *lmx1b* regulation is a feature of more basal gnathostomes, we used a single cell multiomics (RNAseq + ATACseq) approach in the epaulette shark *Hemiscyllium ocellatum*, a representative chondrichthyan and outgroup to osteichthyans. Strikingly, these analyses demonstrated linkages between the *LARM1* enhancer and *lmx1b* in the unpaired dorsal fins of epaulette sharks, as well as the deployment of a Lmx1b-mediated regulatory network whose core downstream components are conserved between dorsal fins, paired fins, and limbs. Collectively, these data support the ancient origin of a “DV” regulatory module in median fins that was redeployed during the early evolution paired fins, facilitating the assembly of the DV patterning axis. Intriguingly, multiome-based inference of global fin network architecture in epaulette shark also revealed reduced regulatory complexity in pectoral relative to median fins, suggesting that co-option as a mechanism for anatomical innovation can proceed through selective streamlining of ancestral network components.

## Introduction

The presence of an appendicular system—whether it be fins for swimming, limbs for running, or wings for flying—is one of the most recognizable features of the jawed vertebrate body plan. Classical hypotheses propose that paired fins first evolved as either transformed gill arches [1], or as retained regions of an elongate ancestral fin that extended along the trunk [2–4]. While new fossil discoveries have invigorated the debate around these hypotheses and shed light on how the vertebrate body plan was assembled (e.g., [5, 6]), the origin of paired fins remains poorly understood. The earliest known vertebrate fossils possessed median fins but lacked paired fins [7–10]. While a variety of paired, lateral fin-like structures were present in jawless vertebrates, the earliest antero-posteriorly restricted paired fins that are often considered homologous to those of jawed vertebrates appeared at the pectoral level in osteostracans, whereas pelvic fins arose later in stem gnathostomes [reviewed in [11–13]; although see [14] for pelvic fin origin of paired fins].

Changes in developmental programs underpin the emergence of novel morphologies. For the origin of the vertebrate paired fins, this transition would have required lateral plate mesodermal cells of the flank to acquire a novel role in producing fin outgrowths [15–18]. Molecular developmental studies on paired fin origins have provided key insights into the evolution of competency zones for appendage formation [19, 20], generative homologies between limbs and gill arches [21–24], the regional patterning of lateral plate mesoderm to form a fin-forming field [15, 18, 25–28], and the early evolution of paired fin musculature [16, 29–34]. Given that median fins predate the evolutionary origin of paired fins, a prominent developmental model for the origin of paired fins posits that pre-existing gene regulatory networks were co-opted from the midline to the flank [35, 36]. Evidence in support of this model includes the similar expression profiles of 5’ *HoxD* and *Tbx* genes in paired and median fins [36], and the deep conservation of Shh-mediated regulatory networks for anterior-posterior patterning and outgrowth [37–40], including the ZRS *Shh* enhancer [41]. Less, however, is known about the evolution of other patterning axes. Here we focus on the evolution of the appendage dorso-ventral (DV) patterning system. While paired fins and limbs have clear top and bottom halves, median fins exhibit left-right mirror symmetry and lack a patterning axis that is topologically equivalent to the DV axis of paired appendages. This raises fundamental questions around how the DV patterning axis arose during vertebrate appendage evolution.

In tetrapods, a core set of molecular interactions pattern limbs along the DV axis [reviewed in [42–44]]. En1, a transcriptional repressor protein present in the ventral ectoderm of limb buds, restricts the expression of the secreted signalling molecule Wnt7a to the dorsal ectoderm [45, 46]. Wnt7a, in turn, induces the expression of Lmx1b, a transcription factor, in the dorsal limb bud mesenchyme, which regulates the formation of dorsal musculoskeletal and integumentary structures, as well as motor axon pathfinding [47–57]. In mice, the loss of *lmx1b* produces a bi-ventralized limb phenotype most apparent in the autopod [42], whereas ectopic expansion of *lmx1b* expression into the ventral limb bud is sufficient to result in double-dorsalized limb morphologies [48, 49, 51, 58, 59]. In mice, Chip-seq experiments have identified two cis regulatory modules, *LARM1* (containing both enhancer and silencer elements) and *LARM2*, that regulate *lmx1b* expression in developing limbs through a positive autoregulatory feedback loop and restrict expression to the dorsal limb compartment [60, 61]. Notably, deletion of *LARM1-LARM2* recapitulates the bi-ventralized limb phenotype of *lmx1b* null mutants without causing negative pleiotropic effects in other tissue types, indicating the limb specificity of the *LARM* enhancer landscape [60].

Less is known about appendage DV patterning systems in finned vertebrates. Dorsoventral asymmetries have been reported in the fin rays (rod like bones that support the distal fin) of the Australian lungfish (*Neoceratodus forsteri*), the non-teleost actinopterygians *Polypterus ornatipinnis* and *Acipenser brevirostrum,* and tetrapodomorph fossils bridging the fin-to-limb transition [62]. In zebrafish, keratinized epidermal structures, termed breeding tubercles, are unique to the dorsal side of the pectoral fins of males and function in reproduction [63, 64]. Molecular studies in zebrafish show that *Wnt7a* and *En1* orthologues compartmentalize the ectoderm along the DV axis, similar to limbs [65–69], and that their expression depends on *fgf10-*mediated signalling [69]. Additionally, *lmx1b* expression has been reported in the dorsally positioned adductor muscles of pectoral fins based on *in situ* hybridization [70]. Because fin myofibres derive from migratory somitic myoblasts [29, 71], these observations contrast with those of limbs, where *lmx1b* is expressed in the lateral plate mesoderm and functions non-cell autonomously in muscle patterning [57, 72]. Among cartilaginous fish, *En1* is expressed in the ventral ectoderm of the pectoral fins of the catshark *Scyliorhinus canicula*, [73], and *lmx1b* in the dorsal mesenchyme of the pectoral and pelvic fins of the little skate *Leucoraja erinacea* [74]. Collectively, these data suggest a *lmx1b*-mediated dorso-ventral patterning system was already in place in the paired fins of the last common ancestor of jawed vertebrates. Most recently, the expression of dorsoventral patterning genes has been characterized in both the paired and unpaired fins of the direct developing cichlid *Astatotilapia burtoni*, the sturgeon *Acipenser baerii,* and the catshark *Scyliorhinus caniculi* [75]. In addition to revealing a limb-like profile of *wnt7a, en1a*, and *lmx1b* in paired fins, these analyses demonstrated broad ectodermal *wnt7a* and posteriorly restricted *lmx1b* expression in the dorsal and anal fins [75]. Interestingly, pharmacological perturbation experiments in *A. burtoni,* provide evidence that Shh- but not Wnt-mediated signalling is required for *lmx1b* expression in the median fins, suggesting the evolution of novel regulatory inputs for dorsal patterning in paired fins [75].

Here we use zebrafish genetics and Epaulette shark multiomics approaches to test predictions of the median fin co-option model with respect to the evolution of the “DV” patterning axis. Using new transgenic tools in zebrafish, we demonstrate that the *lmx1b* lineage contributes to both paired and median fins—forming uniquely regionalized domains that differ between fin types—and that a subset of *lmx1bb-*positive cells differentiates into fin ray osteoblasts. To functionally test the conservation of *lmx1b* regulation between fin types in zebrafish, we use enhancer deletions in *cis* to a *lmx1bb* knock-in fluorescent reporter to show that *LARM* enhancers are principal drivers of *lmx1bb* expression in both the pectoral and pelvic fins but do not contribute to median fin expression. Furthermore, homozygous loss of *LARM1–LARM2* results in a bi-ventralized fin ray phenotype in the paired fins that corresponds to *lmx1bb* lineage contributions but does not affect median fin morphology. To determine if *LARM* regulation of *lmx1b* in paired appendages is a more broadly shared feature of gnathostomes, we generated multiomics data sets for the pectoral and dorsal fins of the Epaulette shark *Hemiscyllium ocellatum* as a representative chondricthyan (sister group to osteichthyans). Remarkably, these analyses identified clear linkages between the LARM regulatory element and *lmx1b* expression in dorsal fins, as well as the deep conservation of a Lmx1b-mediated regulatory network whose core components are conserved between dorsal fins, paired fins, and limbs. Intriguingly, *de-novo* reconstruction of global fin regulatory networks also revealed a greater number of unique connections in median versus paired fins, suggesting network redeployment may be accompanied by reductions in network architecture complexity.

## Results

### *lmx1bb* BAC demonstrates regionalized expression in both the paired and median fins of zebrafish

To characterize the distribution and fate of cells expressing the limb dorsalizing factor *lmx1bb* during fin development in zebrafish, we first generated a stable BAC line [TgBAC(*lmx1bb*:KalTA4_mCherry) in which *lmx1bb* flanking sequence was used to drive an mCherry-KalTA4 cassette (see S1 file for details). Crossing this line with a compound transgenic line that included a KalTA4 responsive mCherry reporter (UAS-Elb:Eco nfsb-mCherry) and ubiquitously expressed GFP (*Ubi*:switch) resulted in embryos with mCherry fluoresence that recapitulated a subset of sites of endogenous *lmx1bb* expression, including the midbrain-hindbrain boundary [76, 77], otic capsules [70, 76], and pectoral fin buds [70] (Figs 1A, 1B, and S1 File). In rendered coronal section, mCherry labelling was restricted to the dorsal compartment of the pectoral fin, demonstrating the BAC promoter was sufficient to organize regionalized fin expression (Fig 1C). As previous reports for mice and zebrafish suggest *lmx1bb* is expressed in different mesodermal lineages [57, 70, 72], we crossed our TgBAC(*lmx1bb*: KalTA4_mCherry) (UAS-Elb:Eco nfsb-mCherry) [hereafter referred to as BAC(*lmx1bb*:mCherry)] line with either a TgBAC(*tbx5a*:GFP) [78, 79] or TgBAC(*pax3a:*GFP) [80, 81] line to label fin-field lateral plate mesoderm or migratory somitic myoblasts, respectively. At 48hpf, *lmx1bb*:mCherry colocalized with *tbx5a*:GFP in cells of the endoskeletal disc and cells surrounding the dorsal muscle anlagen but was absent from *pax3a*:GFP positive myoblasts (Fig 1D–1K), thus confirming their lateral plate identity. To further characterize the expression profile of core components of the limb DV patterning axis in pectoral fins, we performed *in situ* hybridization for zebrafish orthologues of *wnt7a* and *en1*, the upstream canonical regulators of *lmx1b* in limbs. Similar to developing limbs, *wnt7aa* and *en1a* expression was confirmed in the dorsal and ventral pectoral fin ectoderm, respectively (S1 Fig). Together, these observations support the conservation of DV patterning gene spatial domains between teleost paired fins and limbs [65–70, 82].

**Fig 1.**
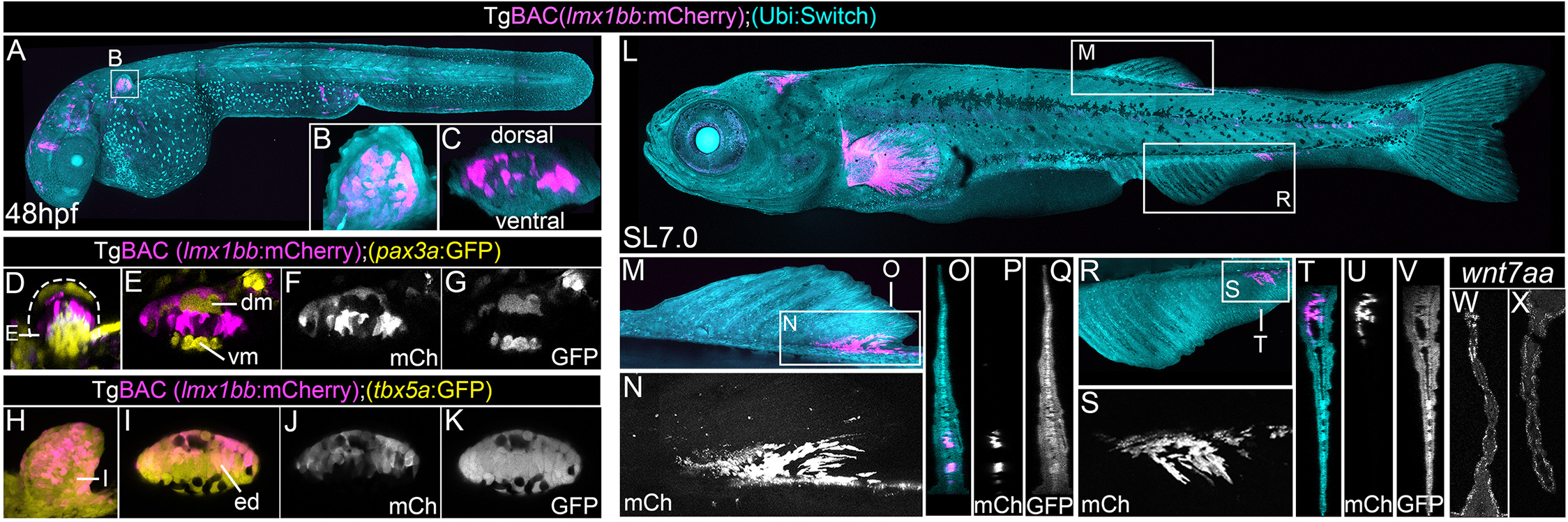
TgBAC(*lmx1bb*:mCherry) transgenic line reveals regionalized *lmx1bb* expression in both paired and median fins of zebrafish. TgBAC(*lmx1bb*: KalTA4_mCherry);(UAS-Elb:Eco-nfsb-mCherry) [hereafter referred to as TgBAC(*lmx1bb*:mCherry)]; (*Ubi*:switch) zebrafish at 48hpf. **(A–C)** Whole embryo (A) and the pectoral fin bud (B) are shown in lateral view. Rendered section of the pectoral fin bud is shown in (C). (**D–G)** Pectoral fin bud of TgBAC(*lmx1bb*:mCherry);(*pax3a*:GFP) at 48hpf shown in lateral view (D) and in rendered section (E–G). *lmx1bb*:mCherry fluorescence does not overlap the somitic myoblast marker *pax3a*:GFP. (**H–K)** Pectoral fin bud of TgBAC(*lmx1bb*:mCherry); (*tbx5a*:GFP) at 48hpf shown in lateral view (H) and in rendered section (I–K). *lmx1bb*:mCherry fluorescence co-localizes with *tbx5a*:GFP in fin lateral plate mesoderm. **(L–V)** TgBAC(*lmx1bb*:mCherry)]; (*Ubi*:switch) zebrafish larvae at standard length (SL) 7.0mm. Whole larvae (L), dorsal fin (M, N) and anal fin (R, S) in lateral view. Rendered sections through the dorsal fin (O–Q) and anal fin (T–V). *lmx1bb*:mCherry fluoresence labels the posterior dorsal and anal fin mesenchyme. **(W, X)** HCR fluorescent in situ hybridization for *wnt7aa* on sections through the dorsal fin (W) and anal fin (X) reveal ectodermal labelling. dm, dorsal muscle mass; GFP, green fluorescent protein; mCh, mCherry; vm, ventral muscle mass.

Given that median fins lack a patterning axis topologically equivalent to the DV axis of limbs, we next characterized BAC(*lmx1bb*:mCherry) expression during median fin development. In zebrafish, two distinct median fins—the dorsal fin and the anal fin—develop from condensations that emerge along a continuous median fin-fold [83–87]. Strikingly, regionalized BAC-*lmx1bb* fluorescence was present in SL 7.0mm larvae at the posterior margin of both the dorsal and anal fins (Fig 1L–1V). Rendered cross-sections of the BAC(*lmx1bb*:mCherry);(*Ubi*:Switch) line confirmed mCherry positive cells were specific to the fin-fold mesenchyme in both median fin types (Fig 1O–1Q, 1T–1V). Given the regionalized expression of *lmx1bb* in median fins, we next sought to determine if other core DV patterning genes were expressed during zebrafish median fin development. Using HCR fluorescent in situ hybridization (FISH), we detected expression of *wnt7aa* along the length of the ectoderm of both the dorsal and anal fins, as well as regions of trunk ectoderm more broadly, but failed to detect *en1* paralogues (Fig 1W, 1X). Together, these results show that zebrafish dorsal and anal fins exhibit the same spatial neighbouring of ectodermal *wnt7aa* and adjacent mesenchymal *lmx1bb* expression posteriorly that characterizes the dorsal compartment of paired appendages.

### *Wnt7a-Lmx1b* expression in median fins is a conserved feature of jawed vertebrates

To determine if the expression of “DV” patterning genes in median fins is a broadly conserved feature of actinopterygians, we next examined the expression of *lmx1b, wnt7a*, and *en1* orthologues in the paddlefish *Polyodon spathula* (Stages 41–43; during fin-fold elaboration) (Fig 2A–2C) as a representative basal (non-teleost) actinopterygian. In both the pectoral and pelvic fins, *wnt7a* and *en1* transcripts were detected in the dorsal and ventral ectoderm, respectively. *lmx1b* expression was present in the dorsal mesenchyme of both fin pairs, but in the pelvic fins it was posteriorly restricted. Like zebrafish, paddlefish possess single dorsal and anal fins [88]. In these median fins, *lmx1b* expression was restricted to the posterior fin mesenchyme and *wnt7a* was expressed more broadly across the fin ectoderm, whereas *en1* was absent.

**Fig 2.**
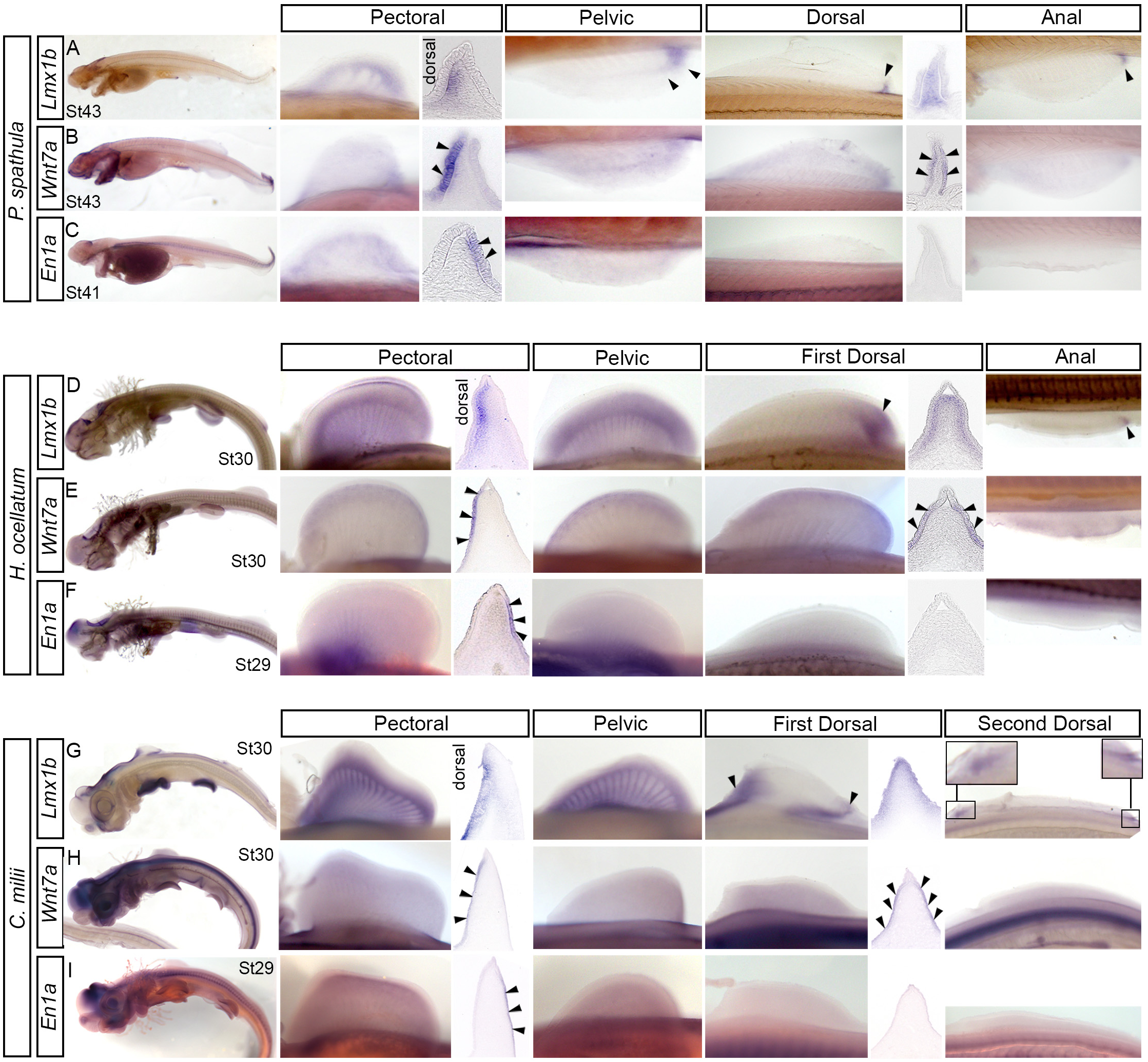
Comparative expression of dorso-ventral patterning genes in paired and median fins. *lmx1b, en1,* and *wnt7a* expression for the paddlefish *Polyodon spathula* **(A–C)**, epaulette shark *Hemiscyllium ocellatum* **(D–F)**, and elephant fish *Callorhinchus milii* **(G–I)**. Whole embryos shown in lateral view, anterior is left. Whole pectoral and pelvic fins are in ventral view, anterior is left. Dorsal and anal fins are in lateral view, anterior is left. For all cross sections, dorsal is left and ventral is right. Arrowheads in sections highlight ectodermal labelling. Arrowheads in whole mounts highlight regionalized expression of *lmx1b*.

To further broaden our comparative framework, we next investigated the expression of “DV” patterning genes in the paired and median fins of epaulette shark (*Hemiscyllium ocellatum*) and elephant fish (*Callorhinchus milii*), representatives of the two major extant lineages of cartilaginous fishes: elasmobranchs (Fig 2D–2F) and holocephalans (Fig 2G–2I), respectively. Both taxa are basal in the gnathostome phylogeny, with *C. milii* (family *Callorhinchidae*) considered to be the most basal of extant holocephalans [89], making it a critical species for examining hypothesis on the origin of paired appendages. In both taxa, the pectoral and pelvic fins exhibited broad expression of *wnt7a* in the dorsal ectoderm, *en1* in the ventral ectoderm, and *lmx1b* in the dorsal mesenchyme. Epaulette sharks possess two morphologically similar dorsal fins and an anal fin. In these unpaired fins, *lmx1b* staining was restricted to the posterior mesenchyme and *wnt7a* was expressed throughout the ectoderm, whereas no expression of *en1* was detected, consistent with the patterns observed in zebrafish and paddlefish. Unlike epaulette sharks, the two dorsal fins of elephant fish differ morphologically. The first dorsal fin is characterized by an ossified spine and a sail-like web, and the second forms as an elongate fin along the tail [90, 91]. Whereas *wnt7a* was expressed along the length of both dorsal fins, *lmx1b* expression was detected in the mesenchyme of both the anterior and posterior domains of each dorsal fin. Collectively, these findings suggest the topological arrangement of *wnt7a*-positive ectoderm adjacent to *lmx1b*-positive mesenchyme in median fins is a shared feature of jawed vertebrates and was likely established prior to the origin of paired fins [75]. Furthermore, the spatial profile of median fin and pelvic fin *lmx1b* expression can vary between lineages, indicating evolutionary flexibility in how *lmx1b*-mediated networks are deployed.

### Regionalized contributions of the Lmx1bb lineage differ between the paired and median fins but include common cell fates

Although *lmx1b* is expressed in mesenchymal cells during both paired and median fin development, it is unknown whether *lmx1b*-expressing cells are maintained throughout fin morphogenesis, which lineages they contribute to, and whether their spatial distribution differs between fin types. To compare *lmx1bb*-expressing progenitor contributions and fates between paired and unpaired fins, we performed a genetic lineage analysis by crossing the BAC(*lmx1bb*:mCherry-KalTA4) line with a Tg(UAS:iCre);(Ubi:switch) compound transgenic line. In resulting progeny, cells with a history of *lmx1bb* BAC-mediated iCre activation undergo a genetic recombination at the *ubi*:Switch locus, resulting in a change from GFP to mCherry fluoresence [92]. Triple transgenic fish were raised to the early juvenile stages (SL8-12mm), at which point the skeletal pattern of the pectoral, dorsal and anal fins is largely established [87, 93, 94]. Tg(UAS:iCre);(*ubi*:Switch) fish were used as a negative control (Fig 3).

**Fig 3.**
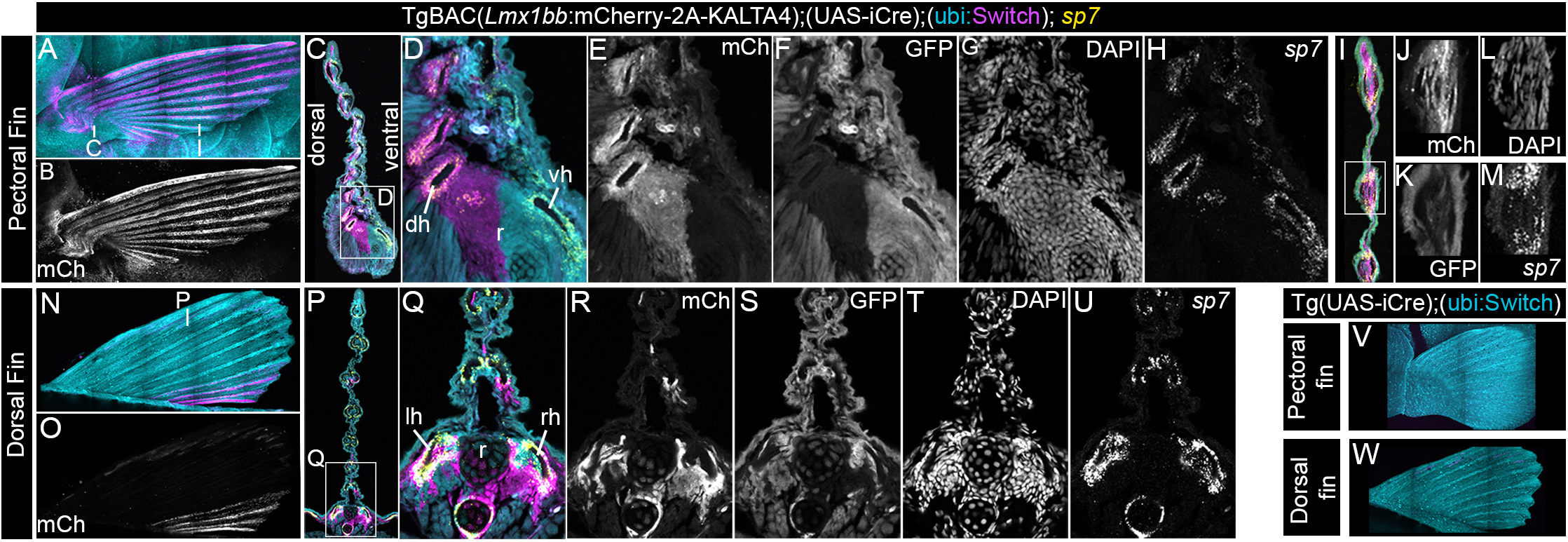
*lmx1bb* genetic lineage tracing in zebrafish pectoral and dorsal fins. Cre-mediated genetic lineage tracing using TgBAC(*lmx1bb*:mCherry-2A-KALTA4);(UAS-iCre);(ubi:Switch). Cells with a history of BAC-*lmx1bb* activation express mCherry, whereas all other cells ubiquitously express GFP. **(A,B)** Whole pectoral fin of zebrafish larvae (standard length, SL = 11 mm) in lateral view; anterior is left. **(C–M)** Cross-sections through the proximal (C–H) and distal (I–M) pectoral fin. Merged channels are shown in (C, D, I). Single channels show mCherry (E, J), GFP (F, K), and DAPI (G, L). HCR fluorescent in situ hybridization for the osteoblast marker *sp7* is shown in (H, M). **(N, O)** Whole dorsal fin of zebrafish larvae (SL = 11 mm) in lateral view; anterior is left. **(P–U)** Cross-section near the posterior margin of the dorsal fin. Merged channels are shown in (P, Q). Single channels show mCherry (R), GFP (S), and DAPI (T). HCR fluorescent in situ hybridization for *sp7* is shown in (U). **(V, W)** Pectoral (V) and dorsal (W) fins of negative-control Tg;(UAS-iCre);(ubi:Switch) zebrafish larvae (SL = 8 mm) in lateral view; anterior is left. mCh, mCherry.

In the pectoral fins of BAC(*lmx1bb*:mCherry-KalTA4);(UAS:iCre);(*ubi*:switch) fish, mCherry labelled cells were visible in the endoskeletal radials and interspersed between myofibres of the adductor muscles (Figs 3A–3G and S2), consistent with their lateral plate derivation. Contributions to the pectoral fin rays (i.e., dermal skeleton) varied along the length of the fin. Proximally, mCherry positive cells surrounded the dorsal half of the fin rays, the dorsal hemitrichia, including along their articulation with the fin endoskeleton (Fig 3C–3G). In contrast, in the distal portion of the rays, mCherry positive cells lined both the dorsal and ventral hemitrichia (Fig 3I–3K, 3M), demonstrating a loss of fidelity of *lmx1bb*-positive cells to the dorsal fin compartment. Given observations that osteoblasts line the hemitrichia of caudal fins [95], we next sought to determine the identity of cells of the *lmx1bb* lineage positioned in close association the dorsal hemitrichia of pectoral fins. Fluorescent in-situ hybridization (FISH) for the osteoblast marker *sp7* confirmed *lmx1bb* patterned cells give rise to osteoblasts of the dorsal but not ventral hemitrichia proximally (Fig 3D, 3E, 3H), but form both dorsally and ventrally positioned osteoblasts distally (Fig 3I–3M).

In dorsal fins, BAC(*lmx1bb*:mCherry-KALTA4) driven iCre resulted in mCherry positive cells in the posterior region of the fin, contributing to the dorsal fin radials as well as to both the left and right hemitrichia of the last several rays (Fig 3N–3T). Similar to the pectoral fins, FISH for *sp7* confirmed that cells of the *lmx1bb* lineage differentiated into osteoblasts of the hemitrichia near their articulation with the endoskeleton (Fig 3Q, 3R, 3U); however, unlike paired fins, this labelling was symmetric across the midline in the unpaired fins. Collectively, these results demonstrate that although the *lmx1bb* lineage contributes to different regions of paired versus median fins, the skeletal derivatives of the *lmx1bb* lineage are similar.

### *LARM cis* regulatory modules are required for zebrafish *lmx1bb* expression in paired but not median fins

The similar topologies of ectodermal *wnt7a* and mesenchymal *lmx1b* expression in both paired and median fins, together with common cell fates derived from the *lmx1bb* lineage, suggest that a pre-existing “DV” appendage regulatory module may have been co-opted from the midline to the flank with the emergence of paired fins. However, the distinct, regionalized domains of *lmx1bb* expression between fin types imply divergent regulatory mechanisms for *lmx1b* induction. In mice, two cis-regulatory modules, *LARM1* and *LARM2,* drive limb *lmx1b* expression, and their loss leads to near-complete abrogation of *lmx1b* transcription and a bi-ventralized phenotype comparable to that of *lmx1b* null mutants. To understand how the *lmx1b* enhancer landscape evolved in vertebrates, we first generated VISTA sequence alignments [96] of regions flanking the *lmx1b* locus (Fig 4A) using a spotted gar viewpoint [97]. Although *LARM1* and *LARM2* have been previously proposed as unique to lobe-finned fish [60], we identified conserved *LARM1* peaks in representative ray-finned and cartilaginous fish, and *LARM2* peaks in ray-finned fishes, suggesting at least a subset of the *LARM* regulatory landscape is conserved across jawed vertebrates.

**Fig 4.**
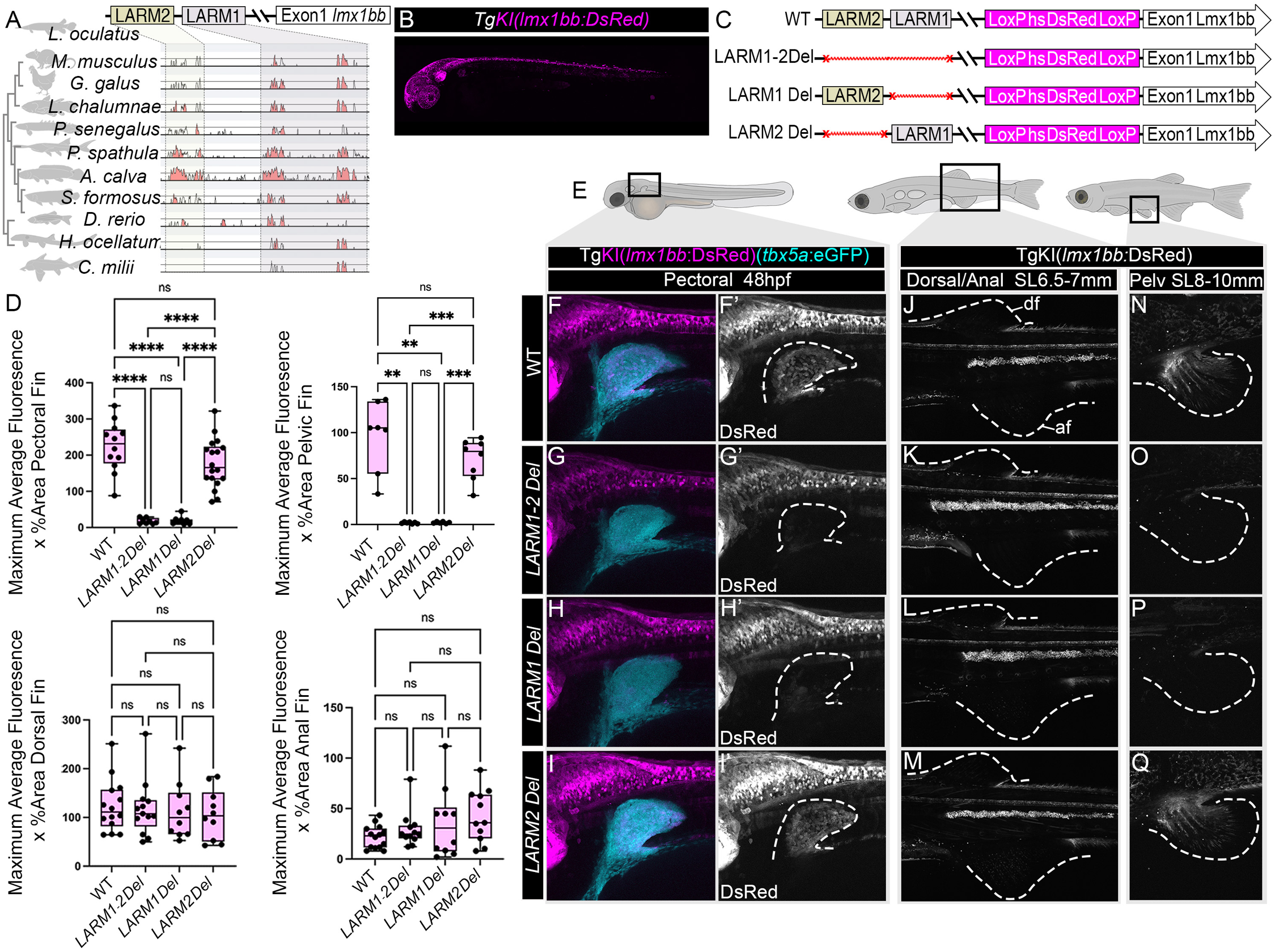
The *LARM1* cis regulatory module is required for paired but not median fin *lmx1b* expression. **(A)** VISTA LAGAN global pairwise sequence alignments of the LARM locus for mouse (*Mus musculus*), chick (*Gallus gallus*), coelacanth (*Latimeria chalumnae*), Senegal bichir (*Polypterus senegalus*), paddlefish (*Polyodon spathula*), bowfin (*Amia calva*), Asian bonytongue (*Scleropages formosus*), zebrafish (*Danio rerio*), epaulette shark (*Hemiscyllium ocellatum*), and elephant fish (*Callorhinchus milii*) with a spotted gar (*Lepisosteus oculatus*) viewpoint. *LARM1* is partially conserved among jawed vertebrates and *LARM2* among bony fishes. **(B)** Tg(*lmx1bb*-LoxP-hsDsRed-LoxP-DTA) knock in line [referred to hereafter as TgKI(*lmx1bb*:DsRed) zebrafish at 48hpf in lateral view, anterior is left. **(C)** LARM deletion series schematic. Single deletions of *LARM1, LARM2*, or a region bracketing *LARM1* + *LARM2* were generated in cis to the TgKI(*lmx1bb*:DsRed) allele. Red line indicates deleted fragment. **(D–Q)** Average maximum fluorescence was compared between WT and *LARM* deletions in cis to the TgKI(*lmx1bb*:DsRed) allele in pectoral (D, F–I’), dorsal, and anal (D, J–M) and pelvic (D, N –Q) fins. All fins are shown in lateral view, anterior is left. The BAC(*tbx5a*:eGFP) transgene is used to visualize the position of the pectoral fin (F–I). In single channel images for DsRed, dashed lines mark fin margins (F’–I’). *LARM1-LARM2* deletions and single *LARM1* deletions resulted in reduced dsRed fluorescence relative to WT for paired but not median fins. Statistical analyses in (D) for pectoral, pelvic and anal fins was done using Welch’s one way ANOVA with Dunnett’s T3 post hoc multiple comparisons test; and for dorsal fins using a one-way ANOVA with Tukey’s multiple-comparisons test. ns is not significant, * is p < 0.05, ** is p < 0.01, *** is p < 0.001, and *** is p < 0.0001.

In order to test the function of *LARM* enhancers in fins, we used CRISPR-Cas9 mediated mutagenesis to generate a series of single-allele germline deletions encompassing regions of sequence conservation for *LARM1*, *LARM2*, or both *LARM1* plus *LARM2* in-cis to a *lmx1bb* knock-in reporter allele [Tg(*lmx1bb*-LoxP-hsDsRed-LoxP-DTA), hereafter referred to as TgKI(*lmx1bb*:DsRed)] (Figs 4B, 4C and S2 file). We then compared fin-specific maximum average fluorescence across mutant and wild type *LARM* backgrounds, providing a direct readout of enhancer function. In both pectoral (48hpf) (Fig 4D–4I’) and pelvic fin buds (SL 8–10mm) (Fig 4D, 4N–4Q), complete deletion of *LARM1*–*LARM2* caused a dramatic reduction in DsRed fluorescence (p<0.0001 for pectoral fins; p<0.01 for pelvic fins). A similar reduction of reporter fluorescence was observed following the singular deletion of *LARM1* (p<0.0001 for pectoral fins; p<0.01 for pelvic fins), whereas no significant reduction in fluorescence was observed following the deletion of the region encompassing *LARM2*. These results demonstrate that *LARM* enhancer activity in paired appendages was already established in the last common ancestor of bony fishes. Furthermore, *LARM* regulatory modules function during the development of both pectoral and pelvic fins, with *LARM1* a principal, paired fin enhancer in zebrafish.

We next sought to test if *LARM* enhancers are required for *lmx1bb* expression in developing dorsal and anal fins, as might be predicted by models of median fin regulatory network co-option during the origin of paired fins. In contrast to pectoral and pelvic fins, complete deletions of *LARM1–LARM2* or single deletions of *LARM1* or *LARM2* did not significantly affect dsRed fluorescence in the dorsal or anal fins (Fig 4D, 4J–4M). Together, these results provide evidence that distinct *LARM-*dependent and independent regulatory systems operate in zebrafish paired versus median fins, respectively.

Given the role of *wnt7a* as an upstream regulator of *lmx1b* in limbs, and the expression of *wnt7aa* in the pectoral [69], dorsal, and anal fins of zebrafish larvae (Fig 1W, 1X), we generated a stable *wnt7aa* mutant with a premature stop in the second exon (S3A Fig), which was then back-crossed with our TgKI(*lmx1bb*:DsRed) knock-in allele. Consistent with a unique regulation, no differences in *lmx1b* driven DsRed fluorescence was observed in the dorsal fins of WT fish compared to *wnt7aa* heterozygous or homozygous mutants (S3B and S3C Fig), though we cannot exclude the possibility of compensation by *wnt7ab* paralogues.

### Deletion of *LARM* enhancers affects zebrafish paired fin but not median fin patterning

To determine if *LARM* enhancers are selectively required for patterning of paired versus median fins in zebrafish, we generated a germline *LARM1–LARM2* deletion mutant in a WT background (S2 File). Progeny of in-crossed heterozygotes were raised to late sub-adult stages (approximately 1.7–2.4cm SL), scanned by micro-computed tomography (uCT), and digital reconstructions used to assess skeletal pattern. In the pectoral fins of wild type (n=4) and heterozygous (n=8) zebrafish, the basal portion of the dorsal and ventral hemitrichia were asymmetric across the midline, such that the dorsal hemitrichia formed a relatively smooth border near their articulation with the endoskeleton, whereas the ventral hemitrichia formed a prominent saw-tooth pattern when visualized in lateral and medial view (Fig 5A–5D). Strikingly, in all *LARM1–LARM2* null mutants scanned (n=5), the pattern of the ventral hemitrichia was duplicated in the dorsal fin compartment, resulting in a bi-ventralized phenotype (Fig 5E–5H).

**Fig 5.**
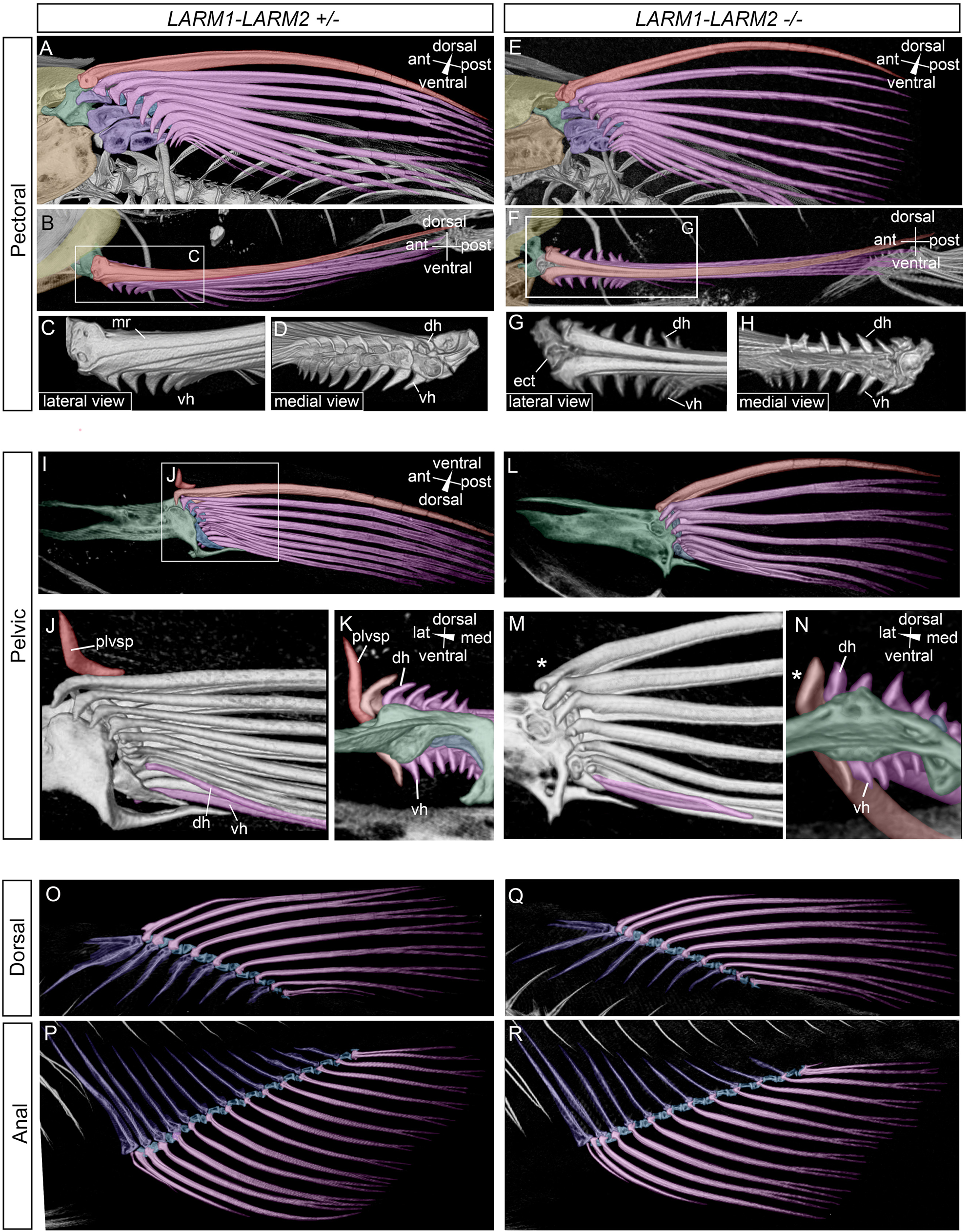
*LARM* deletion results in aberrant skeletal patterning of the paired fins but not median fins. Digital renderings of the fin skeleton from microcomputed tomography scans. **(A–H)** Pectoral fins of *LARM1–LARM2*^+/-^ heterozygotes (A–D) and *LARM1-LARM2*^-/-^homozygous (E–H) mutants. In (A, B, E, F), the skeletal elements of the shoulder girdle and pectoral fin are pseudocolored as follows: cleithrum, yellow; coracoid, brown; scapula, green; proximal radials, blue; distal radials, cyan; marginal ray, pink and all other rays magenta. Radials are partially removed in (C) and (D). The dorsal hemitrichia (dh) of the fin rays appear bi-ventralized in *LARM1–LARM2^-/-^* mutants and form a sawtooth pattern similar to the ventral hemitrichia (vh). (**I–N)** Pelvic fins of *LARM1-LARM2*^+/-^ heterozygotes (I–K) and *LARM1-LARM2*^-/-^ homozygous (L–N) mutants. In (I, K, L, N) the pelvic girdle is pseudocolored green; radials, blue; the marginal ray, pink; and all other rays magenta. In (J), a single ray is pseudocolored magenta to highlight lateral curvature of the dorsal hemitrichia, which forms a “Y” shape with the ventral hemitrichia in *LARM1-LARM2*^+/-^ heterozygotes. In *LARM1-LARM2*^-/-^ mutants, this curvature is lost, the dorsal hemitrichia appear ventralized, and the pelvic splint (plvsp, red) is absent (*). **O–R**. No differences were observed in the dorsal (O, Q) or anal (P, R) fins between LARM1-LARM2^+/-^ heterozygotes (O, P) and LARM1-LARM2^-/-^ homozygous (Q, R) mutants. The proximal radials are pseudocolored blue; the distal radials, cyan; and the fin rays, magenta. ect, ectopic bone; mr, marginal ray.

Like the pectoral fins, loss of *LARM1–LARM2* also resulted in dorsal duplications of ventral morphologies in the pelvic fins. In wild type and heterozygous fish, the dorsal hemitrichia curve laterally after splitting from the ventral hemitrichia, creating a deep Y-shape that is most pronounced for the medial rays (Figs 5I, 5J, S4A and S4B). In contrast, in homozygous mutants, the dorsal hemitrichia remain in register with the ventral hemitrichia, resulting in a narrower “Y” (Figs 5L, 5M, and S4C-S4E, p<0.01). Furthermore, the shape of the proximal ends of the dorsal hemitrichia appear ventralized in homozygous mutants relative to heterozygotes (Fig 5N). We also identified a prominent, dorsally directed spine lateral to the first fin ray in all wild type (n=4) and a subset of heterozygous (50%, n=4/8) fish examined by microCT. To the best of our knowledge, this element has not been reported in zebrafish but appears similar to the pelvic splint reported in a variety of teleost fishes [98–100]. This bone was absent in all homozygous mutants (n=0/5), indicating its formation is *LARM-*enhancer dependent (S4F-S4I Fig, p<0.01).

In addition to the pectoral fins, we examined the dorsal and anal fins of *LARM1/LARM2*+/+, *LARM1/LARM2*+/-, and *LARM1/LARM2*-/- fish. We did not observe any difference in the number of rays or skeletal morphologies (Fig 5O–5R). Collectively, these data are consistent with our enhancer-deletion knock-in reporter assays (Fig 4) and demonstrate that the *LARM* regulatory module is essential for dorsoventral patterning of the paired fin dermal skeleton but is not required for patterning of the median fins in zebrafish.

Because *LARM1-LARM2* deletion removes most but not all reporter expression (Fig 4), we sought to compare enhancer deletion phenotypes with those of *lmx1bb* null mutants. Using CRISPR-Cas9 gene editing, we generated a stable germ line *lmx1bb* mutant with a premature stop in the third exon (S5A Fig). However, homozygous mutants were embryonic lethal, precluding analyses of skeletal phenotypes (S5B Fig).

### Multiomics analysis of epaulette shark fins supports *LARM1* regulatory linkage and Lmx1b-mediated network co-option from dorsal fins

To investigate the evolutionary origins of *lmx1b* regulatory modules, we used 10X Genomics single-nucleus multiome sequencing (snRNA-seq + snATAC-seq) to profile pectoral and dorsal fins from Stage 30–31 epaulette shark (*Hemiscyllium ocellatum*) embryos as a representative chondrichthyan. To enable these analyses, we recently generated a high-quality, fully annotated, trio-based reference genome for *H. ocellatum* [101]. Resulting data sets were processed in Seurat (v5.3.0) and Signac (v1.15.0), with dimensionality reduction and clustering performed on both RNA and ATAC modalities. In total, 2,891 nuclei from the pectoral fins and 1,765 nuclei from dorsal fins were recovered. Using combined transcriptomic and ATAC data sets, we identified 14 molecularly distinct clusters, which were annotated based on marker gene profiles from published limb datasets [102–105] (Fig 6A). Clusters included ectoderm, chondrogenic, myogenic, connective tissue, neural crest and dermal skeleton cell populations. Clusters without clear cell type affinities were annotated based on gene enrichment (see Table in S4 File for cluster-specific gene enrichments). *lmx1b* was expressed in a subset of cells across multiple clusters, including those enriched for the dermal skeleton marker *And1* and molecularly diverse mesenchymal/connective tissue populations (Fig 6B, 6C).

**Fig 6.**
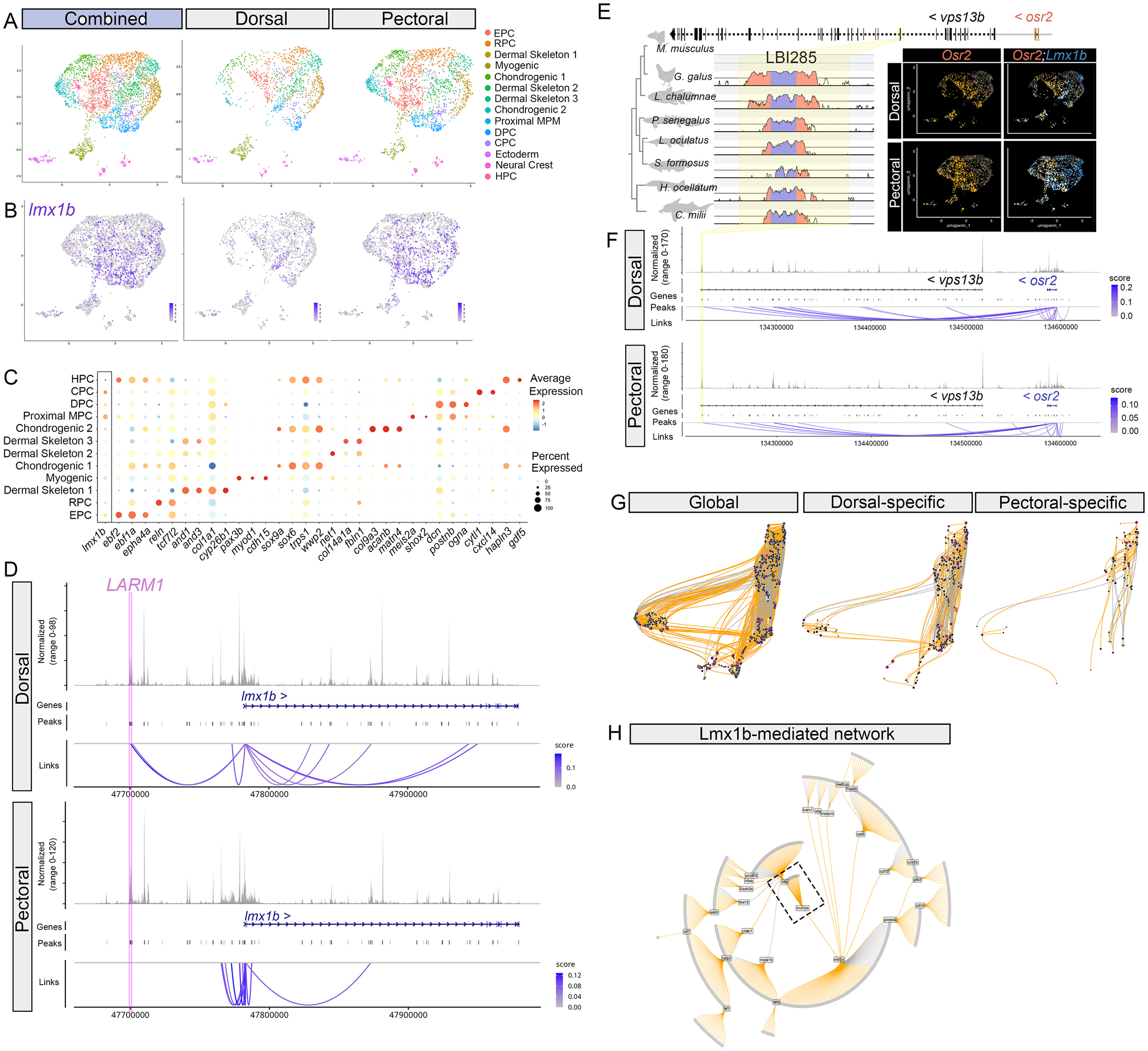
Pectoral and dorsal fin multiome of epaulette shark. **(A)** UMAP visualization of cell clusters in St.30/31 *H. ocellatum* pectoral and dorsal fins based on single nuclei RNA and ATAC sequencing. Cell identities are color coded and based on limb marker gene expression or gene enrichment. **(B)** UMAP visualization of *lmx1b* expression across clusters. **(C)** Dot plot showing *lmx1b* and select marker gene expression across clusters. **(D)** Link-Peak anlaysis (Signac) of *lmx1b* locus. Regulatory linkages are shown in purple and represent correlations between peak accessibility and gene expression. The epaulette shark orthologue of the mouse *LARM1* element (magenta) is linked with *lmx1b* in dorsal fins. **(E)** Schematic of mouse genomic locus containing the direct Lmx1b binding enhancer *LBI285* and its regulatory target *osr2* (top row). VISTA alignment of *LBI285* in chick (*Gallus gallus*), coelacanth (*Latimeria chalumnae*), Senegal bichir (*Polypterus senegalus*), spotted gar (*Lepisosteus oculatus*), Asian bonytongue (*Scleropages formosus*), epaulette shark (*Hemiscyllium ocellatum*), and elephant fish (*Callorhinchus milii*). The *LBI285* enhancer spans across an exon (blue) within *vps13b*. Conserved sequence is pink. UMAP visualization of *lmx1b* (blue) and *osr2* (orange) expression, which colocalizes in a subset of cells in each fin type. **(F)** Link-Peak analysis (Signac) of *osr2*. The position of the Epaulette *LBI285* orthologue is highlighted with yellow bar and is linked to *osr2* expression in both pectoral and dorsal fins. **(G)** UMAP visualization global fin regulatory network (left) generated with PANDO, as well as dorsal fin specific (middle) and pectoral fin specific (right) networks following pruning of shared edges. Orange lines indicate positive linkages; gray lines indicate negative linkages. Circles are nodes and color represents centrality in network. **(H)** Recalculated plot of Lmx1b-mediated network. Dashed box highlights position of *lmx1b* and its proximate downstream targets in network. EPC, *ebf2*-positive cells; RPC, *reln*-positive cells; MPC, *meis2a*-positive cells; DPC, *decorin*-positive connective tissue; CPC, *cytl1*-postiive cells; HPC, *hapln3*-positive cells.

Given that *LARM* enhancer function is required for limb [60] and zebrafish paired-fin patterning, but is dispensable for zebrafish dorsal and anal fins, we next asked whether *LARM* regulates *lmx1b* expression in epaulette shark paired versus median fins. To address this, we used peak–gene linkage analysis across an approximate 300kb region bracketing the Lmx1b start site (LinkPeaks, Signac) [106, 107] to identify ATAC peaks whose accessibility covaries with *lmx1b* transcription (Fig 6D). This analysis identified seven dorsal fin–specific peak–gene linkages, eight pectoral fin–specific linkages, and a single shared peak near the proximal promoter. Importantly, we detected a linkage between the *LARM1* enhancer and *lmx1b* in dorsal fins, contrary to predictions based on zebrafish. To test the robustness of this association, we used Cicero to assess peak-peak co-accessibility and generate a regulatory map of the *lmx1b* locus [108]. This analysis similarly supported a linkage between *LARM1* and the *lmx1b* promoter in the dorsal fin (S6 Fig). Together, these epaulette shark results, though correlative, support conserved *LARM*-mediated regulation of *lmx1b* expression in the dorsal fin within a broader and more complex fin enhancer landscape. Although a comparable linkage between *LARM1* and *lmx1b* in the pectoral fins of epaulette sharks was not captured in our St.30/31 data set, this may reflect stage-specific differences in the dynamics of *lmx1b* regulation, as limb enhancer function can vary temporally [109].

To test if assembly of the paired fin dorso–ventral patterning axis involved redeployment of a median fin Lmx1b-mediated network, we used both a candidate-gene approach and *de novo* network inference in epaulette shark. In mice, combined ChIP-seq [61] and mutant transcriptomic analyses [52, 55, 110] have defined core components of an Lmx1b-mediated dorsal determination program [61]. To assess the potential antiquity of Lmx1b “limb” networks in shark paired and median fins, we first used VISTA alignments to test whether mouse Lmx1b-binding elements are sequence-conserved across finned vertebrates. We identified conserved shark orthologues for 22 of the 292 reported mouse elements, 15 of which maintained synteny within 1Mb of their projected target genes (Table in S5 File). Multiome profiling showed that, in both pectoral and dorsal fins, these conserved elements overlapped accessible ATAC peaks and that their associated target genes were co-expressed with *lmx1b* (Figs 6E and S7; Table in S5 File), consistent with predictions of gene network co-option. Conserved mouse-shark *lmx1b* modules included genes implicated in limb connective tissue, skeleton, and joint formation (e.g., *Shox2* [111–114], *Nfia* [115, 116], Osr2 [117, 118], Trps1 [119], Ankrd11 [120]), as well as myogenesis (Tcf12: [121]), suggesting ancestral roles in musculoskeletal and ECM development. To further test whether enhancer accessibility covaried with the transcription of predicted target genes, we performed peak-to-gene linkage analysis [106, 107], which recovered positive associations for a subset of these genes (S7 Fig). The linkages for Osr2 provide one such example (Fig 6E, 6F).

Recently developed bioinformatic pipelines enable inference of gene regulatory network architecture from multiomics data sets [122, 123]. In addition to evaluating candidate pathways based on known limb Lmx1b targets, we used Pando [124] to model fin gene regulatory networks by integrating chromatin accessibility, transcript abundance, and predicted transcription factor binding motif sites. We first generated an initial global fin network, which was then pruned to edges that were differentially accessible in each fin type (Fig 6G) [124]. Interestingly, despite similar profiles in ATAC and RNAseq quality and coverage between fin types (S8 Fig), and a slightly greater peak count in pectoral fins (pectoral fin n_peaks=262,354; dorsal fin n_peaks=230,803), pectoral fins showed reduced unique pathway complexity relative to dorsal fins (Fig 6G). We next recalculated the network graph to identify Lmx1b-associated modules, recovering 34 predicted direct Lmx1b targets (Fig 6H). Remarkably, seven of these 34 inferred genes were orthologues of, or closely related family members to, direct Lmx1b targets reported in mice (Table in S6 File) [61]. In direct support of predictions of gene network co-option, most inferred direct Lmx1b-target modules (24/34) were shared between pectoral and dorsal fins, whereas the remainder were fin type specific (Table in S6 File). These observations from both candidate-based analyses and de-novo network inference, support a model in which core components of an ancient Lmx1b-mediated regulatory network were already in place in median fins, and the evolutionary acquisition of *lmx1b* expression in incipient paired fins enabled redeployment of these modules to facilitate assembly of the dorso-ventral patterning axis. Moreover, while overall similarity between dorsal and pectoral fin total network architecture is consistent with co-option during early fin evolution, the smaller number of fin-specific connections in the pectoral fin suggests a reduction in network complexity accompanied this redeployment.

## Discussion

The evolutionary acquisition of paired fins created a new dorsoventral anatomical axis. Defining the cellular and molecular bases for establishing dorsal versus ventral morphologies is therefore central to understanding early fin evolution. We have mapped the contributions of the *lmx1bb* lineage to mature pectoral and dorsal fin anatomy in zebrafish, revealing common chondrocyte, connective tissue, and dermal skeleton osteoblast fates across fin types. These findings broadly align with *lmx1b* Cre-based lineage tracings in mouse limbs, which similarly identified chondrocyte and connective tissue derivatives [57, 125]. Fin ray reduction and eventual loss in the tetrapod lineage [106] would have required loss of a dermal skeletal fate from the *lmx1b*-lineage tree, potentially occurring through the erosion of direct Lmx1b-responsive cis-regulatory targets. However, because fin rays persist in *LARM* deletion mutants, it is unlikely that loss of Lmx1b-mediated pathways would have been a primary driver of dermal skeleton loss. Instead, such changes were likely integrated within a broader suite of developmental mechanisms for dermal skeletal reduction and endoskeletal elaboration [126–133].

In zebrafish, deletion of *LARM* regulatory modules causes a deficit in *lmx1b* expression in the pectoral and pelvic fins and results in fin-ray patterning defects. Strikingly, the biventralized phenotype of the pectoral fin dorsal hemitrichia closely correlates with the distribution of *lmx1b* lineage-derived osteoblasts. A recent bioRxiv preprint by Hawkins et al [134] similarly used zebrafish mutants to dissect LARM enhancer function during paired and median fin development and, through cross-species transgenic assays, showed that *LARM* elements from diverse gnathostomes, including skate and catshark, can drive fluorescent reporter expression in zebrafish pectoral fins. Our *LARM* deletion phenotypes broadly agree with those of Hawkins et al. [134], together supporting deep conservation of an *lmx1b* dorsal determination program in paired fins that is regulated by a *LARM* cis-regulatory module. Although our epaulette shark multiome peak-linkage analysis did not recover a linkage between *LARM1* and *lmx1b* in the pectoral fins, the ability of skate and catshark *LARM1* enhancers to drive reporter expression in zebrafish pectoral fins further supports the antiquity of enhancer function across paired fins to the base of gnathostomes [134].

Interestingly, Hawkins et al. [134] also reported that whereas limb-like Wnt7a-Engrailed-Lmx1b epistasis is maintained in early zebrafish pectoral fins, *en1a; en1b* double mutant zebrafish do not exhibit an overt pectoral fin phenotype, as would be predicted with ventral expansion of *lmx1b* expression [49, 135]. In mice, the bi-dorsalized phenotype of *En1* mutants predominantly affects the autopod (i.e, distal-most region of limb) [49, 136, 137]. Our Cre-mediated fate mapping in zebrafish shows that dorsal-specific contributions of *lmx1bb*-expressing cells are not maintained in distal fin rays where both dorsal and ventral hemitrichia derive from a *lmx1bb*-positive lineage. Given evidence for generative homologies between the autopod and fin rays [126], this loss of dorsal lineage fidelity may help explain the absence of an *en1* mutant fin phenotype in fins.

Comparative expression analyses support the conservation of ectodermal *wnt7a* and mesenchymal *lmx1b* expression domains in the unpaired fins of gnathostomes [75, 134]. Although shared expression patterns can suggest regulatory network co-option, such similarities can also arise convergently. Demonstrating the same cis-regulatory element drives expression of a network component in two different developmental contexts provides a key test of co-option models [138]. The long-range ZRS enhancer for *Shh* is one such example, functioning in both limbs and unpaired fins [41]. In zebrafish, however, deletion of the *LARM* cis-regulatory module affects pectoral and pelvic fins but not median fins (see also Hawkins et al. [134]), and *wnt7aa* is dispensable for *lmx1bb* expression. Together, these observations for zebrafish run counter to predictions of simple gene network co-option [134].

Chondrichthyans, as the sister group to osteichthyans, occupy a critical phylogenetic position for reconstructing the early evolution of vertebrate fin regulatory systems [37, 73, 127, 132, 133, 139–147]. Our multiomics-based analysis in epaulette shark identified both distinct, fin-specific sets of predicted cis-regulatory elements at the *lmx1b* locus, as well as a clear linkage between *LARM1* and *lmx1b* activation in dorsal fins, supporting the ancient origin of *LARM* function in unpaired fins and its reuse in paired fins. Furthermore, our candidate-based analyses of direct Lmx1b targets, as well as our de novo assembly of fin regulatory networks, define an ancient Lmx1b-mediated core network shared between shark dorsal fins and mouse limbs. How this median fin Lmx1b-module was effectively “rotated” 90° in paired fins and integrated with flank-derived regulatory inputs is not well understood, but likely occurred in the context of a primitive body wall ectoderm already compartmentalized along the DV axis by *engrailed* [73, 75, 148]. Under such a co-option model, the absence of an effect of *LARM* deletion on *lmx1b* expression in zebrafish dorsal and anal fins may reflect *lmx1b* enhancer redundancy, consistent with our identification of multiple predicted dorsal-fin regulatory elements in shark. Alternatively, lineage specific differences in median fin *LARM* deployment may have evolved between cartilaginous fish and teleosts (or ray-finned fish more broadly).

Beyond DV patterning, our Pando-based assembly of global fin networks in Epaulette shark was consistent with broad scale network redeployment, as most inferred nodes and edges were shared across paired and median fins. Intriguingly, however, dorsal fins exhibited greater fin-specific pathway complexity than pectoral fins. Because paired fins were secondarily added to an ancestral axial system, the comparatively simpler network architecture of pectoral fins may reflect redeployment of only subsets of an ancestral median fin network (i.e., partial co-option, [149]), potentially related to differences in embryonic context [150–152]. Alternatively, an initially broad co-option from median to paired fins may have been followed by network refinement, as has been proposed for butterfly eyespots [153]. Though speculative, differences in the degree of enhancer redundancy may provide one molecular correlate of this broader disparity in network complexity and could potentially account for the impact of *LARM* deletion on paired but not median fins. Future work to characterize the *lmx1b* regulatory landscape will be critical to evaluating this hypothesis.

Collectively, the studies we describe here reveal the signature of an ancient “DV” regulatory module in median fins that was redeployed during the early evolution of paired fins, facilitating the assembly of the DV patterning axis and dorsal determination. In doing so, this addresses a key limitation of simple co-option models by providing a molecular mechanistic framework for how a DV axis, absent in median fins, could have evolved in paired fins. Interestingly, while it is possible that the first paired fins were symmetric structures, redeployment of an ancestral median fin core *lmx1b* network raises the possibility that DV patterning asymmetries may have been intrinsic features of incipient paired fins from their origin. Given the range of Lmx1b-mediated pathways, such asymmetries could have involved patterning of the skeleton and musculature, ECM organization, or motor axon pathfinding (see also ref [75]).

Our analyses of broader network architectures also reveal that the emergence of paired fins may have been accompanied by a reduction in the regulatory elements associated with unpaired fins. This suggests that increasing morphological complexity may not be necessarily driven by the expansion of regulatory and enhancer landscapes, but instead, the evolution of novel structures may, in some cases, involve strategic simplification or removal of existing regulatory networks. How widely a “less is more” mechanism for regulatory network re-purposing applies across vertebrate phylogeny remains unknown but is now amenable to systematic investigation through the expanding application of comparative genomic technologies across an increasingly broad range of taxa.

## Methods

### Animal handling and transgenic lines

Adult zebrafish were maintained in a recirculating freshwater aquarium system kept at approximately 28°C with a 14-hour light, 10-hour dark cycle using standard husbandry procedures [154]. Zebrafish work was done in accordance with Monash Animal Ethics Committee approved project IDs 22161, 25827, and 41803. Previously described lines included TgBAC(*pax3*:GFP)^i150^ [81], Tg (*tbx5a*:eGFP) [78, 79]; and Tg(*ubi*:switch) [92]. The Tg(*lmx1bb*-LoxP-hsDsRed-LoxP-DTA) line was obtained from the National BioResource Project, Zebrafish (https://shigen.nig.ac.jp/zebra/). Embryos of the paddlefish *Polyodon spathula* were obtained from Osage Catfisheries Inc. (Osage Beach, MO, USA), raised in 18°C freshwater tanks at Kennesaw State University (Kennesaw, GA, USA) as previously described [127] and in accordance with KSU Institutional Animal Care and Use Committee approved protocols 12-001, 16-001. Embryos of the Epaulette shark *Hemiscyllium ocellatum* were obtained from an adult brood stock maintained at Monash University (Clayton, VIC, AUS), using procedures for Epaulette husbandry, breeding, and egg collection as previously described [101] and in accordance with protocols approved under Monash University Animal Ethics Project IDs 30347 and 44673. Embryos of the Elephant fish *Callorhincus milii* were obtained from wild caught adults temporarily maintained in captivity for egg laying using previously described capture and husbandry protocols [155] in accordance with Monash University Animal Ethics Project IDs 22254 and 40651. Staging of Paddlefish, Epaulette shark, and fish were based on references [88, 90, 156, 157].

To generate the TgBAC(*lmx1bb*: KalTA4_mCherry) strain, the BAC clone (see S1 File for details) was obtained from the BACPAC resource centre (https://bacpacresources.org) and modified by the Monash Genome Modification platform to generate the final Lmx1bb BAC clone used for injection. In brief, a KalTA4-T2A-mCherry-polyA-FRT-neomycin-FRT cassette was inserted and FLP recombination was used to remove the neomycin resistance cassette. Finally, an iTOL2 - KM cassette was added and the BAC simultaneously trimmed down to a total length of approximately 103 Kb. The final BAC was injected at 100 ng/μl into one-cell-stage wildtype embryos along with transposase RNA (25 ng/μl) synthesized from the pcs2FA-transposase vector using an mMessage machine Sp6 kit (Ambion, AM1340). Injected embryos were manually sorted for mCherry fluorescence. Positive embryos were raised to adults and outcrossed for founders to establish a stable line.

CRISPR/Cas9-mediated gene editing was used to generate mutants used in this study. crRNA sequences were designed using either the IDT (https://sg.idtdna.com/site/order/designtool/index/CRISPR_CUSTOM) or ChopChop (https://chopchop.cbu.uib.no) [158] online tools. *LARM1, LARM2,* and *LARM1-LARM2* mutants were generated by targeted deletion using pairs of crRNA sequences listed in S3 file. Targeting crRNAs were duplexed with tracrRNA according to the manufacturer’s protocol to generate gRNAs, which were subsequently injected along with Cas9 protein (Alt-R CRISPR–Cas9 system, IDT) into one-cell-stage heterozygous [Tg(*lmx1bb*-LoxP-hsDsRed-LoxP-DTA) embryos for LARM deletion mutants. Stable germline mutants for LARM enhancers *in cis* to the (*lmx1bb*-LoxP-hsDsRed-LoxP-DTA) transgene included a net deletion of 2682bp for *LARM1*; a deletion of 764bp for *LARM2*; and 6827bp deletion for the region spanning *LARM1-LARM2*. The stable germline mutant for *LARM1-LARM2* deletion in a wild type background (used for assessing skeletal malformations) included a 6862bp deletion. Mutants for *wnt7aa* and *lmx1bb* were generated by insertion of a premature STOP cassette at cut sites generated using the crRNA sequences listed in S3 File. STOP cassette oligonucleotide design and co-injection was performed as previously described [159]. Oligonucleotide sequences included a universal STOP cassette and flanking gene-specific homology arms, as listed in S3 File. Primers used for genotyping of mutant lines are listed in S3 File.

### cDNA synthesis, cloning and in situ hybridization

Cloning for ISH probe template was done as previously described [133]. RNA was isolated from pooled embryos with either Trizol reagent (Invitrogen) plus the PureLink^TM^ Micro-to-Midi Total RNA Purification System (Invitrogen) for *P. spathula*, or with a RNAeasy Mini Kit (Qiagen) for *D. rerio, H. ocellatum, and C. milii,* and was used as template for synthesizing single-strand cDNA with the Superscript III First Strand Synthesis kit (Invitrogen), all per manufacturer’s instructions. Primer pairs for cloning probe fragments are listed in S3 File.

PCR amplicons were cloned into pGEM®-T Easy Vector (Promega) and verified by Sanger sequencing. For *in situ* hybridization, Plasmids were linearized with restriction enzymes (New England Biolabs, NEB). Any 3′ overhangs were blunted using DNA Polymerase I, Large Klenow (NEB). In vitro transcription was carried out using SP6 or T7 RNA polymerases (Promega) and Digoxigenin RNA labeling mixes (Roche) according to manufacturer′s instructions. *In situ* hybridizations were as described in Modrell et al. [160]. Hybridization was carried out at 68–70 °C overnight. Digoxigenin-labelled probes were detected with Anti-Digoxigenin-AP, Fab fragments (1:2000 dilution; Roche) and developed with BM-Purple (Roche). Sense probe was used with stage-matched specimens as a negative control.

### Hybridization Chain Reaction

Hybridization Chain reaction RNA FISH probe against zebrafish *sp7* and *wnt7aa* were generated using a custom HCR Probe Designer [161] and included 27 probe pairs designed for use with the B3 amplifier for *sp7*; and 24 probe pairs designed for use with the B1 amplifier for *wnt7aa*. Probe sequences are presented in the table in File S7. FISH was performed on slides according to manufacturer’s recommendations (https://www.molecularinstruments.com). Each probe set was used at a concentration of 0.4 pmol per 100uL of hybridization buffer, and hybridization was carried out overnight at 37°C in a humidity chamber. H1 and H2 hairpins were used at a concentration of 6pmols per 100 microliters of amplification buffer, and amplification conducted for approximately 18 hours at room temperature. Sections were imaged on a Zeiss LSM 980 Confocal Microscope.

### Fluorescence assays in TgKI(*lmx1bb*:DsRed)] background

All wild type control and *LARM* deletion mutants in the TgKI(*lmx1bb*:DsRed) background, as well as *wnt7aa* mutants in the TgKI(*lmx1bb*:DsRed)] background, were imaged using a Zeiss 980LSM confocal microscope and Zen Blue edition software. Image acquisition settings were identical across wild type and mutant fish for each fin type. Image stacks were analyzed using Fiji (ImageJ) software. For LARM deletion analyses, a baseline minimum threshold for pixel intensity was set to ten for pectoral and pelvic fins and to 20 for dorsal and anal fins. For Wnt7aa mutant analyses, a baseline minimum threshold for pixel intensity was set to ten. Fins were traced in each Z-stack section, the mean intensity multiplied by % area of fluorescence within each fin tracing, and the maximum value used for each fish.

### VISTA alignments

Genome sequence data was obtained from Ensembl (https://asia.ensembl.org/index.html) or the NIH Genome database (https://www.ncbi.nlm.nih.gov/gdv). Conserved sequence elements were identified using mVISTA LAGAN (https://genome.lbl.gov/vista/mvista/submit.shtml) with translated anchoring to improve alignments of distal homologies, a 20bp calc window, 40bps min cons width, and a 70% conserved identity threshold.

### Micro-computed tomography

Zebrafish used for X-ray tomography were fixed overnight in 4% PFA, washed in PBS and mounted in plastic tubes in 1% agarose for scanning. Scans were carried out at the Monash Monash Department of Civil Engineering on a Zeiss Xradia 520 Versa and image sets visualized using Avizo Software (Thermo Fischer Scientific).

### Isolation of nuclei from epaulette shark fins

Three individual stage 30–31 epaulette shark embryos were euthanized in 0.1g/L tricaine methanesulphonate solution and the pectoral and dorsal fins dissected into 1ml of cell isolation buffer [1 x Hank’s Balanced Salt Solution (HBSS) containing 5% Fetal Bovine Serum (FBS)] on ice. Once fin dissections were complete, samples were incubated at 32°C for 30 minutes in 1ml of collagenase solution (1 x HBSS, 5% FBS, 1% Collagenase type II) with regular mixing. Samples were pelleted by centrifugation at 2500rpm at 4°C for 5 minutes, resuspended in 1 ml cell isolation buffer, and passed through a 70um cell strainer (Corning) followed by a 40um cell strainer (Corning). Samples were then pelleted by centrifugation at 2500rpm at 4°C for 5 minutes, resuspended in 1ml cell isolation buffer and stained with Propidium Iodide. Live cells were fluorescence sorted using an Influx cell sorter by the FlowCore platform (Monash University) and transported to PeterMac for downstream multiome processing.

For nuclei isolation, live cells were fixed with formaldehyde to preserve nuclear integrity. Formaldehyde was added to the cell suspension to a final concentration of 1%, followed by gentle inversion to mix. Samples were incubated at room temperature for 10 min and quenched with 0.125 M glycine. Cells were washed once with 1 mL ice-cold PBS, inverted 5 times to mix, and centrifuged at 500 × g for 5 min at 4 °C. The supernatant was discarded, and the pellet was resuspended in 1 mL ice-cold PBS.

The cell pellet then was resuspended in 100 µL chilled Low Loss Lysis (LLL) buffer containing (10 mM Tris-HCl, 10 mM NaCl, 3 mM MgCl₂, 0.1% NP-40, 1% BSA, 1 mM DTT, 0.2 U/µL RNase inhibitor) and incubated on ice for 3 min to gently lyse cell membranes. Subsequently, 1 mL of chilled Wash Buffer containing (10 mM Tris-HCl, 10 mM NaCl, 3 mM MgCl₂, 1% BSA, 1 mM DTT, 0.2 U/µL RNase inhibitor) was added, and the sample was centrifuged again at 500 × g for 5 min at 4 °C. The supernatant was discarded, and the nuclei pellet was resuspended in 1× Nuclei Buffer prepared from 20× Nuclei Buffer supplemented with 1 mM DTT and 0.5 U/µL RNase inhibitor. The nuclei suspension was then passed through a 40 µm Flowmi® strainer into a low-binding 1.5 mL Eppendorf tube.

### 10x Genomics multiome library preparation and sequencing

The nuclei were diluted to a final concentration that enabled a recovery of 10,000. The Chromium Next GEM single Cell Multiome ATAC+ Gene Expression reagent (PN-1000283) was used. The nuclei were tagmented using ATAC buffer B and ATAC enzyme B and incubated at 37 °C for 60 minutes. Barcoding master mix was then added to the tagmented nuclei. We used Chip J and Multiome reagents to run 10X Chromium to generate GEMs (following the 10xUser Guide CG000338 Rev F). GEMs were cleaned up and libraries were generated according to the 10x User Guide CG000338 manual.

Gene expression libraries were sequenced on the NovaSeq 6000 (paired- end 150 bp reads, targeting 30,000-40,000 reads per cell). ATAC libraries were sequenced on the NovaSeq 6000 (paired- end 150 bp reads, targeting 25,000 reads per cell).

### Multiome and gene regulatory network analysis

The multiomic dataset was processed using cellranger-arc (v 2.0.2) with default parameters. cellranger-arc mkref was first run using a custom reference generated using the NCBI epaulette shark genome (GCF_020745735.1). The annotation file was filtered to keep only protein-coding genes and lncRNA. Two annotations were manually added to the reference (And1, And3). LOC132825693 was renamed to Lmx1bb for downstream compatibility with Pando/Jaspar reference.

After running cellranger-arc, MACS3 was run individually on the atac_fragments.tsv.gz file to generate a new sample peak set (-f BED --nomodel --keep-dup all -g 3.9e+09 --shift -37 --extsize 73). The peak sets were then merged together into a consensus peak set. The data was then analysed using Seurat (v.5.3.0) & Signac (1.15.0). The multiomic data was loaded individually and FeatureMatrix() was used to quantify reads against the consensus MACS3 peak set. Standard Seurat/Signac control checks were performed and low quality cells were removed. The two samples were merged together. Dimensionality reduction and clustering were performed on both the RNA and ATAC datasets. The ATAC data was batch corrected using harmony (1.2.3). FindMultiModalNeighbors was used to perform joint analysis of the RNA and ATAC modalities.

AddMotifs was run with the Jaspar2020 core vertebrate motifs to find motif sequence information and FindMotifs was used to find enriched motifs in the dataset. A motif-tf-gene table was generated by querying biomaRt based on the enriched motifs. Pando was run using the motif table and the Seurat object to perform gene regulatory network analysis. find_modules was run with lax parameters (p_thresh = 0.1, nvar_thresh = 2, min_genes_per_module = 1, rsq_thresh = 0.05).

Cicero was run separately on the dorsal and pectoral datasets. Both datasets were processed independently following the standard Cicero workflow. Peaks with zero counts were removed and a UMAP was calculated. run_cicero was used with a subset of genes and default parameters.

### Statistical analysis

GraphPad Prism was used to analyse data sets. All graphs show the number of fish used in each experiment. For pectoral, pelvic and anal fins, maximum average fluorescence between wild type, *LARM1-LARM2, LARM1*, and *LARM2* deletion mutants were compared using Welch’s one way ANOVA and Dunnett’s T3 multiple-comparisons test. For dorsal fins, assumptions of equal variances (Brown-Forsythe) were met, and one-way ANOVA performed with Tukey’s multiple-comparisons test. Fin ray widths below the split of the dorsal and ventral hemitrichia were compared using a Kruskal–Wallis test with Dunn’s multiple-comparisons test (adjusted p values; α = 0.05). All other fin ray measurements were compared using a one-way ANOVA with Tukey’s multiple-comparisons test. p values less than 0.05 were considered significant.

## Supporting information

## Supplementary Files

**S1 File.** Generation of TgBAC(*lmx1bb:*mcherry) and comparison with endogenous expression.

**S2 File.** *LARM* regulatory module deletion.

**S3 File.** Table of CRISPR guide sequences, genotyping primers, and *in situ* hybridization cloning primers.

**S4 File.** Table of cluster gene enrichments for St.30/31 *H. ocellatum* pectoral and dorsal fins based on single nuclei RNA and ATAC sequencing.

**S5 File.** Table of conservation of mouse Lmx1b binding elements in epaulette shark

**S6 File.** Table of Pando inferred Lmx1b direct regulatory targets in St.30/31 epaulette shark dorsal and pectoral fins.

**S7 File.** Table of Fluorescent In situ hybridization probe sets for zebrafish

## Supplementary Figure Legends

**S1 Fig. Expression of *wnt7aa* and *en1a* in 48hpf zebrafish**. In situ hybridization for *en1a* **(A, B)** and *wnt7aa* **(C, D)** in 48hpf zebrafish embryos. Whole mounts are shown in lateral view, anterior is left (A, C). In section (B, D), dorsal is right, and ventral is left.

**S2 Fig. *Lmx1bb* lineage analysis in zebrafish pectoral fins.** Genetic lineage analysis using the TgBAC(*lmx1bb*:mCherry-2A-KALTA4);(UAS-iCre);(ubi:Switch). Cells with a history of BAC-*lmx1bb* activation express mCherry (magenta), whereas all other cells ubiquitously express GFP (cyan). Actin staining (yellow) with phalloidin makes visible myofibers, and DAPI counterstain labels nuclei. **(A)** Cross section through the proximal region of the pectoral fin of larvae (Standard length, SL 11mm). **(B–K)** Enlarged insets from (A) showing merged (B, G) and individual channels (C–F, H–K). mCherry positive cells contribute to fin radials (r), dorsal hemitrichia (dh), and surrounding connective tissues. Arrowheads (H) highlight subset of mCherry positive cells in adductor muscle (ad). ab, abductor muscle; dh, dorsal hemitrichia; r, radials; vh, ventral hemitrichia.

**S3 Fig. TgKI(*lmx1bb*:DsRed) fluorescence in the dorsal fin is not affected in *wnt7aa* mutants.** (**A)** CRISPR/Cas9-mediated introduction of a premature STOP codon (red asterisk) in the second exon of *wnt7aa*. Pink box indicates mutated sequence, which was confirmed by Nanopore sequencing. **(B, C)** Comparison of maximum average fluorescence of TgKI(*lmx1bb*:DsRed) knock-in allele between *wnt7aa*+/+, *wnt7aa*+/-, and *wnt7aa*-/- mutants. In (C), dorsal fins are shown in lateral view, anterior is left. No difference in maximum average fluorescence was observed. Maximum average dorsal fin fluorescence was compared using a one-way ANOVA with Tukey’s multiple-comparisons test. ns, not significant (p>0.05).

**S4 Fig. *LARM1-LARM2* deletion results in bi-ventralized skeletal patterning of the pelvic fins and loss of pelvic splint. (A–C)** Digital renderings of the pelvic fin skeleton from microcomputed tomography scans for *LARM1-LARM2*^+/+^ (A), *LARM1-LARM2^+/-^* (B) and *LARM1-LARM2^-/-^* (C) fish. Segmentation-based reconstruction of the medial-most fin ray is shown in blue both overlain on the fin (A–C) and in isolation (A’–C’). (**D, E)** Fin ray schematic (D) indicating positions of measurements used for quantifications (E). No differences between fin ray width below split of the dorsal and ventral hemitrichia (“lower” width) were observed among genotypes (Kruskal–Wallis with Dunn’s post hoc, all adjusted p > 0.05). Neither the width of separation between the distal margins of the dorsal (dh) and ventral hemitrichia (vh) (upper width), nor the upper/lower width ratios, differed between *LARM1-LARM2*+/+ and *LARM1-LARM2*+/-genotypes. However, both the upper width and the ratio of upper/lower width were significantly different in LARM1-LARM2-/- (one-way ANOVA with Tukey’s multiple-comparisons test, **, p<0.001 and *** p< 0.0001). **(F–I)** Digital renderings of the partial pelvic fin skeleton from microcomputed tomography scans for *LARM1–LARM2^+/+^* (F), *LARM1–LARM2^+/-^* (G, H) and *LARM1–LARM2^-/-^* (I) fish. * indicates the absence of the pelvic splint (plvsp). **(J)** Pelvic splint (plvsp) presence and absence was compared across genotypes using a Fisher’s exact test. Splint presence differed by genotype (p=0.0105); pairwise testing showed WT differed from homozygous mutants (**, p=0.0079), while WT vs heterozygotes (p=0.2081) and heterozygotes vs homozygotes (p = 0.1049) were not significant.

**S5 Fig. *lmx1bb* homozygous null mutants are embryonic lethal. (A)** CRISPR/Cas9-mediated introduction of a premature STOP codon (red *) in the third exon of *lmx1bb*. Sanger sequencing confirmed mutation. **(B)** Dorsal and lateral views of 7dpf *lmx1bb+/+, lmx1bb+/-, lmx1bb-/-* zebrafish. Scale bar=500un.

**S6 Fig. Peak-peak linkage analysis in epaulette shark.** Cicero-based cis-regulatory map of an approximately 100kb interval at the Lmx1b locus from single cell ATAC profiles of St.30/31 epaulette shark dorsal (**A**) and pectoral fins (**B**). Y-axis indicates co-accessibility. Pink box indicates position of the *LARM1* element. Gray box indicates position of proximal promoter.

**S7 Fig. Multiome link-peak analyses of epaulette shark orthologues of Lmx1b-regulated limb targets. (A, D, G)** Top: Schematic of mouse genomic locus for mouse Lmx1b-binding cis regulatory element *LBI481* and its target gene *shox2* (A); *LBI373* and its target gene *tcf4* (D); and *LBI392* and its target gene *zdhhc6* (G). Bottom: VISTA alignments for *LBI481* (A) and *LBI373* (D) include chick (*Gallus gallus*), coelacanth (*Latimeria chalumnae*), Senegal bichir (*Polypterus senegalus*), spotted gar (*Lepisosteus oculatus*), Asian bonytongue (*Scleropages formosus*), epaulette shark (*Hemiscyllium ocellatum*), and elephant fish (*Callorhinchus milii*). VISTA alignment for *LBI392* (G) include chick (*Gallus gallus*), coelacanth (*Latimeria chalumnae*), Senegal bichir (*Polypterus senegalus*), spotted gar (*Lepisosteus oculatus*), epaulette shark (*Hemiscyllium ocellatum*), and elephant fish (*Callorhinchus milii*). All alignments are from a mouse viewpoint, and regions of conserved sequence are pink. **(B, E, H)** UMAP visualization of *lmx1b* (blue) and either *shox2* (B), *tcf4* (E) or *zdhhc6* (H) (orange) expression based on single nuclei RNA and ATAC sequencing of St.30/31 *H. ocellatum* pectoral and dorsal fins. Double positive cells are present in both pectoral and dorsal fins for all three putative target genes. **(C, F, I)** Link-Peak analysis (Signac) of *shox2* (C), *tcf4* (F), and *zdhhc6* (I) for dorsal fins. Linkages between ATAC peak accessibility and gene transcription are shown in purple. The approximate positions of *LBI* cis regulatory elements are highlighted in yellow. The conservation of mouse Lmx1b-binding cis regulatory elements in sharks, together with their positive regulatory linkages to predicted target genes, supports deep conservation of Lmx1b-mediated regulatory networks between shark dorsal fins and limbs.

**S8 Fig. Quality metrics for St.30/31 Epaulette shark multiome data sets for pectoral and dorsal fins.** RNA-seq QC metrics include RNA nFeatures (A) and RNA nCount (B). ATAC-seq QC metrics include ATAC nFeatures (C); ATAC nCount (D); Peak enrichment at transcription start sites (TSS) (E); nucleosome signal (F); and fraction of reads in peaks (FRiP).

## Acknowledgements

We thank the National BioResource Project, Zebrafish, and Dr. Shin-ichi Higashijima, National Institute of Natural Science, Japan, for providing the (*Imxlbb*-hs:loxP-DRed-loxP-DTA) zebrafish strain, Monash Micro Imaging for imaging assistance, Monash FlowCore for sorting of cells using FACS, and Monash Fishcore staff for assistance with zebrafish and shark husbandry.

## References

1. Gegenbaur C. Zur morphologie der gliedmassen der wirbelthiere. Morph Jahrb. 1876;2:396.

2. Thacher JK. Median and paired fins, a contribution to the history of vertebrate limbs. Trans Conn Acad Sci. 1877;3:281.

3. Mivart SG. XII. Notes on the fins of ‘Elasmobranchs, with Considerations on the Nature and Homologues of Vertebrate Limbs. The Transactions of the Zoological Society of London. 1879;10(10):439–84.

4. Balfour FM, editor On the Development of the Skeleton of the Paired Fins of Elasmobranchii, considered in Relation to its Bearings on the Nature of the Limbs of the Vertebrata. Proceedings of the Zoological Society of London; 1881: Wiley Online Library.

5. Gai Z, Li Q, Ferrón HG, Keating JN, Wang J, Donoghue PC, et al. Galeaspid anatomy and the origin of vertebrate paired appendages. Nature. 2022;609(7929):959–63. doi: 10.1038/s41586-022-04897-6.

6. Brazeau MD, Castiello M, El Fassi El Fehri A, Hamilton L, Ivanov AO, Johanson Z, et al. Fossil evidence for a pharyngeal origin of the vertebrate pectoral girdle. Nature. 2023;623(7987):550–4. doi: 10.1038/s41586-023-06702-4.

7. Shu D. A paleontological perspective of vertebrate origin. Chinese Science Bulletin. 2003;48(8):725–35. doi: 10.1007/BF03187041.

8. Shu DG, Luo H-L, Conway Morris S, Zhang XL, Hu S-X, Chen L, et al. Lower Cambrian vertebrates from south China. Nature. 1999;402(6757):42–6.

9. Xian-guang H, Aldridge RJ, Siveter DJ, Siveter DJ, Xiang-hong F. New evidence on the anatomy and phylogeny of the earliest vertebrates. Proceedings of the Royal Society B: Biological Sciences. 2002;269(1503):1865–9. doi: 10.1098/rspb.2002.2104.

10. Zhang XG, Hou XG. Evidence for a single median fin-fold and tail in the Lower Cambrian vertebrate, Haikouichthys ercaicunensis. Journal of Evolutionary Biology. 2004;17(5):1162–6. doi: 10.1111/j.1420-9101.2004.00741.x.

11. Coates MI. The origin of vertebrate limbs. Development. 1994;1994(Supplement):169–80. doi: 10.1242/dev.1994.Supplement.169.

12. Coates MI. The evolution of paired fins. Theory in Biosciences. 2003;122(2):266–87. doi: 10.1007/s12064-003-0057-4.

13. Larouche O, Zelditch ML, Cloutier R. A critical appraisal of appendage disparity and homology in fishes. Fish and Fisheries. 2019;20(6):1138–75.

14. Wilson M, Hanke G, Märss T. Paired fins of jawless vertebrates and their homologies across the “agnathan”-gnathostome transition. Major transitions in vertebrate evolution. 2007:122–49.

15. Onimaru K, Shoguchi E, Kuratani S, Tanaka M. Development and evolution of the lateral plate mesoderm: Comparative analysis of amphioxus and lamprey with implications for the acquisition of paired fins. Developmental Biology. 2011;359(1):124–36. doi: 10.1016/j.ydbio.2011.08.003.

16. Tulenko FJ, McCauley DW, MacKenzie EL, Mazan S, Kuratani S, Sugahara F, et al. Body wall development in lamprey and a new perspective on the origin of vertebrate paired fins. Proceedings of the National Academy of Sciences. 2013;110(29):11899–904. doi: 10.1073/pnas.1304210110.

17. Onai T. Organization of the body wall in lampreys informs the evolution of the vertebrate paired appendages. Journal of Morphology. 2023;284(3):e21559. doi: 10.1002/jmor.21559.

18. Tzung K-W, Lalonde RL, Prummel KD, Mahabaleshwar H, Moran HR, Stundl J, et al. A median fin derived from the lateral plate mesoderm and the origin of paired fins. Nature. 2023;618(7965):543–9. doi: 10.1038/s41586-023-06100-w.

19. Yonei-Tamura S, Abe G, Tanaka Y, Anno H, Noro M, Ide H, et al. Competent stripes for diverse positions of limbs/fins in gnathostome embryos. Evolution & Development. 2008;10(6):737–45. doi: 10.1111/j.1525-142X.2008.00288.x.

20. Sleight VA, Gillis JA. Embryonic origin and serial homology of gill arches and paired fins in the skate, Leucoraja erinacea. eLife. 2020;9:e60635. doi: 10.7554/eLife.60635.

21. Gillis JA, Dahn RD, Shubin NH. Shared developmental mechanisms pattern the vertebrate gill arch and paired fin skeletons. Proceedings of the National Academy of Sciences. 2009;106(14):5720–4. doi: doi:10.1073/pnas.0810959106.

22. Gillis JA, Hall BK. A shared role for sonic hedgehog signalling in patterning chondrichthyan gill arch appendages and tetrapod limbs. Development. 2016;143(8):1313–7. doi: 10.1242/dev.133884.

23. Gillis JA, Rawlinson KA, Bell J, Lyon WS, Baker CV, Shubin NH. Holocephalan embryos provide evidence for gill arch appendage reduction and opercular evolution in cartilaginous fishes. Proceedings of the National Academy of Sciences. 2011;108(4):1507–12. doi: doi:10.1073/pnas.1012968108.

24. Rees JM, Sleight VA, Clark SJ, Nakamura T, Gillis JA. Ectodermal Wnt signaling, cell fate determination, and polarity of the skate gill arch skeleton. Elife. 2023;12:e79964. doi: 10.7554/eLife.79964.

25. Minguillon C, Gibson-Brown JJ, Logan MP. Tbx4/5 gene duplication and the origin of vertebrate paired appendages. Proceedings of the National Academy of Sciences. 2009;106(51):21726–30. doi: 10.1073/pnas.0910153106.

26. Tanaka M, Onimaru K. Acquisition of the paired fins: a view from the sequential evolution of the lateral plate mesoderm. Evolution & Development. 2012;14(5):412–20. doi: 10.1111/j.1525-142X.2012.00561.x.

27. Adachi N, Robinson M, Goolsbee A, Shubin NH. Regulatory evolution of Tbx5 and the origin of paired appendages. Proceedings of the National Academy of Sciences. 2016;113(36):10115–20. doi: 10.1073/pnas.1609997113.

28. Tanaka M. Fins into limbs: Autopod acquisition and anterior elements reduction by modifying gene networks involving 5’Hox, Gli3, and Shh. Developmental Biology. 2016;413(1):1–7. doi: 10.1016/j.ydbio.2016.03.007.

29. Neyt C, Jagla K, Thisse C, Thisse B, Haines L, Currie P. Evolutionary origins of vertebrate appendicular muscle. Nature. 2000;408(6808):82–6. doi: 10.1038/35040549.

30. Okamoto E, Kusakabe R, Kuraku S, Hyodo S, Robert-Moreno A, Onimaru K, et al. Migratory appendicular muscles precursor cells in the common ancestor to all vertebrates. Nature Ecology & Evolution. 2017;1(11):1731–6.

31. Kusakabe R, Tanaka M, Kuratani S. Developmental Evolution of Hypaxial Muscles: Insights From Cyclostomes and Chondrichthyans. Frontiers in Cell and Developmental Biology. 2021;9:760366. doi: 10.3389/fcell.2021.760366.

32. Ziermann JM, Freitas R, Diogo R. Muscle development in the shark Scyliorhinus canicula: implications for the evolution of the gnathostome head and paired appendage musculature. Frontiers in zoology. 2017;14(1):31.

33. Adachi N, Pascual-Anaya J, Hirai T, Higuchi S, Kuroda S, Kuratani S. Stepwise participation of HGF/MET signaling in the development of migratory muscle precursors during vertebrate evolution. Zoological letters. 2018;4(1):18. doi: 10.1186/s40851-018-0094-y.

34. Turner N, Mikalauskaite D, Barone K, Flaherty K, Senevirathne G, Adachi N, et al. The evolutionary origins and diversity of the neuromuscular system of paired appendages in batoids. Proceedings of the Royal Society B: Biological Sciences. 2019;286(1914). doi: 10.1098/rspb.2019.1571.

35. Freitas R, Gómez-Skarmeta JL, Rodrigues PN. New frontiers in the evolution of fin development. Journal of Experimental Zoology Part B: Molecular and Developmental Evolution. 2014;322(7):540–52. doi: 10.1002/jez.b.22563.

36. Freitas R, Zhang G, Cohn MJ. Evidence that mechanisms of fin development evolved in the midline of early vertebrates. Nature. 2006;442(7106):1033–7. doi: 10.1038/nature04984.

37. Dahn RD, Davis MC, Pappano WN, Shubin NH. Sonic hedgehog function in chondrichthyan fins and the evolution of appendage patterning. Nature. 2007;445(7125):311–4. doi: 10.1038/nature05436.

38. Höch R, Schneider RF, Kickuth A, Meyer A, Woltering JM. Spiny and soft-rayed fin domains in acanthomorph fish are established through a BMP-gremlin-shh signaling network. Proceedings of the National Academy of Sciences. 2021;118(29):e2101783118. doi: 10.1073/pnas.2101783118.

39. Hawkins MB, Jandzik D, Tulenko FJ, Cass AN, Nakamura T, Shubin NH, et al. An Fgf–Shh positive feedback loop drives growth in developing unpaired fins. Proceedings of the National Academy of Sciences. 2022;119(10):e2120150119. doi: 10.1073/pnas.2120150119.

40. Letelier J, Naranjo S, Sospedra-Arrufat I, Martinez-Morales JR, Lopez-Rios J, Shubin N, et al. The Shh/Gli3 gene regulatory network precedes the origin of paired fins and reveals the deep homology between distal fins and digits. Proceedings of the National Academy of Sciences. 2021;118(46):e2100575118. doi: 10.1073/pnas.2100575118.

41. Letelier J, de la Calle-Mustienes E, Pieretti J, Naranjo S, Maeso I, Nakamura T, et al. A conserved Shh cis-regulatory module highlights a common developmental origin of unpaired and paired fins. Nature Genetics. 2018;50(4):504–9. doi: 10.1038/s41588-018-0080-5.

42. Chen H, Johnson RL. Dorsoventral patterning of the vertebrate limb: a process governed by multiple events. Cell and Tissue Research. 1999;296(1):67–73. doi: 10.1007/s004410051267.

43. McQueen C, Towers M. Establishing the pattern of the vertebrate limb. Development. 2020;147(17). doi: 10.1242/dev.177956.

44. Castilla-Ibeas A, Zdral S, Oberg KC, Ros MA. The limb dorsoventral axis: Lmx1b’s role in development, pathology, evolution, and regeneration. Developmental Dynamics. 2024;253(9):798–814. doi: 10.1002/dvdy.695.

45. Rodriguez-Esteban C, Schwabe JWR, Peña JDL, Foys B, Eshelman B, Belmonte JCI. Radical fringe positions the apical ectodermal ridge at the dorsoventral boundary of the vertebrate limb. Nature. 1997;386(6623):360–6. doi: 10.1038/386360a0.

46. Laufer E, Dahn R, Orozco OE, Yeo C-Y, Pisenti J, Henrique D, et al. Expression of Radical fringe in limb-bud ectoderm regulates apical ectodermal ridge formation. Nature. 1997;386(6623):366–73. doi: 10.1038/386366a0.

47. Parr BA, McMahon AP. Dorsalizing signal Wnt-7a required for normal polarity of D–V and A–P axes of mouse limb. Nature. 1995;374(6520):350–3. doi: 10.1038/374350a0.

48. Riddle RD, Ensini M, Nelson C, Tsuchida T, Jessell TM, Tabin C. Induction of the LIM homeobox gene Lmx1 by WNT6a establishes dorsoventral pattern in the vertebrate limb. Cell. 1995;83(4):631–40.

49. Loomis CA, Harris E, Michaud J, Wurst W, Hanks M, Joyner AL. The mouse Engrailed-1 gene and ventral limb patterning. Nature. 1996;382(6589):360–3. doi: 10.1038/382360a0.

50. Loomis CA, Kimmel RA, Tong C-X, Michaud J, Joyner AL. Analysis of the genetic pathway leading to formation of ectopic apical ectodermal ridges in mouse Engrailed-1 mutant limbs. Development. 1998;125(6):1137–48. doi: 10.1242/dev.125.6.1137.

51. Cygan JA, Johnson RL, McMahon AP. Novel regulatory interactions revealed by studies of murine limb pattern in Wnt-7a and En-1 mutants. Development. 1997;124(24):5021–32. doi: 10.1242/dev.124.24.5021.

52. Krawchuk D, Kania A. Identification of genes controlled by LMX1B in the developing mouse limb bud. Developmental Dynamics. 2008;237(4):1183–92. doi: 10.1002/dvdy.21514.

53. Kania A, Jessell TM. Topographic Motor Projections in the Limb Imposed by LIM Homeodomain Protein Regulation of Ephrin-A:EphA Interactions. Neuron. 2003;38(4):581–96. doi: 10.1016/S0896-6273(03)00292-7.

54. Kania A, Johnson RL, Jessell TM. Coordinate Roles for LIM Homeobox Genes in Directing the Dorsoventral Trajectory of Motor Axons in the Vertebrate Limb. Cell. 2000;102(2):161–73. doi: 10.1016/S0092-8674(00)00022-2.

55. Feenstra JM, Kanaya K, Pira CU, Hoffman SE, Eppey RJ, Oberg KC. Detection of genes regulated by Lmx1b during limb dorsalization. Development, Growth & Differentiation. 2012;54(4):451–62. doi: 10.1111/j.1440-169X.2012.01331.x.

56. Havis E, Coumailleau P, Bonnet A, Bismuth K, Bonnin M-A, Johnson R, et al. Sim2 prevents entry into the myogenic program by repressing MyoD transcription during limb embryonic myogenesis. Development. 2012;139(11):1910–20. doi: 10.1242/dev.072561.

57. Li Y, Qiu Q, Watson SS, Schweitzer R, Johnson RL. Uncoupling skeletal and connective tissue patterning: conditional deletion in cartilage progenitors reveals cell-autonomous requirements for Lmx1b in dorsal-ventral limb patterning. Development. 2010;137(7):1181–8. doi: 10.1242/dev.045237.

58. Chen H, Johnson RL. Interactions between dorsal-ventral patterning genes lmx1b, engrailed-1 and wnt-7a in the vertebrate limb. International Journal of Developmental Biology. 2002;46(7):937–42.

59. Vogel A, Rodriguez C, Warnken W, Belmonte JCI. Dorsal cell fate specified by chick Lmxl during vertebrate limb development. Nature. 1995;378(6558):716–20. doi: 10.1038/378716a0.

60. Haro E, Petit F, Pira CU, Spady CD, Lucas-Toca S, Yorozuya LI, et al. Identification of limb-specific Lmx1b auto-regulatory modules with Nail-patella syndrome pathogenicity. Nature Communications. 2021;12(1):5533. doi: 10.1038/s41467-021-25844-5.

61. Haro E, Watson BA, Feenstra JM, Tegeler L, Pira CU, Mohan S, et al. Lmx1b-targeted cis-regulatory modules involved in limb dorsalization. Development. 2017;144(11):2009–20. doi: 10.1242/dev.146332.

62. Stewart TA, Lemberg JB, Taft NK, Yoo I, Daeschler EB, Shubin NH. Fin ray patterns at the fin-to-limb transition. Proceedings of the National Academy of Sciences. 2020;117(3):1612–20. doi: 10.1073/pnas.1915983117.

63. McMillan SC, Géraudie J, Akimenko M-A. Pectoral Fin Breeding Tubercle Clusters: A Method to Determine Zebrafish Sex. Zebrafish. 2015;12(1):121–3. doi: 10.1089/zeb.2014.1060.

64. McMillan SC, Xu ZT, Zhang J, Teh C, Korzh V, Trudeau VL, et al. Regeneration of breeding tubercles on zebrafish pectoral fins requires androgens and two waves of revascularization. Development. 2013;140(21):4323–34. doi: 10.1242/dev.095992.

65. Ekker M, Wegner J, Andrée Akimenko M, Westerfield M. Coordinate embryonic expression of three zebrafish engrailed genes. Development. 1992;116(4):1001–10. doi: 10.1242/dev.116.4.1001.

66. Grandel H, Draper BW, Schulte-Merker S. dackel acts in the ectoderm of the zebrafish pectoral fin bud to maintain AER signaling. Development. 2000;127(19):4169–78.

67. Hatta K, Bremiller R, Westerfield M, Kimmel CB. Diversity of expression of engrailed-like antigens in zebrafish. Development. 1991;112(3):821–32. doi: 10.1242/dev.112.3.821.

68. Neumann CJ, Grandel H, Gaffield W, Schulte-Merker S, Nüsslein-Volhard C. Transient establishment of anteroposterior polarity in the zebrafish pectoral fin bud in the absence of sonic hedgehog activity. Development. 1999;126(21):4817–26. doi: 10.1242/dev.126.21.4817.

69. Norton WHJ, Ledin J, Grandel H, Neumann CJ. HSPG synthesis by zebrafish Ext2 and Extl3 is required for Fgf10 signalling during limb development. Development. 2005;132(22):4963–73. doi: 10.1242/dev.02084.

70. Uemura O, Okada Y, Ando H, Guedj M, Higashijima S-i, Shimazaki T, et al. Comparative functional genomics revealed conservation and diversification of three enhancers of the isl1 gene for motor and sensory neuron-specific expression. Developmental Biology. 2005;278(2):587–606. doi: 10.1016/j.ydbio.2004.11.031.

71. Cole NJ, Hall TE, Don EK, Berger S, Boisvert CA, Neyt C, et al. Development and Evolution of the Muscles of the Pelvic Fin. PLOS Biology. 2011;9(10):e1001168. doi: 10.1371/journal.pbio.1001168.

72. Schweizer H, Johnson R, Brand-Saberi B. Characterization of migration behavior of myogenic precursor cells in the limb bud with respect to Lmx1b expression. Anatomy and Embryology. 2004;208(1):7–18. doi: 10.1007/s00429-003-0373-y.

73. Tanaka M, Münsterberg A, Anderson WG, Prescott AR, Hazon N, Tickle C. Fin development in a cartilaginous fish and the origin of vertebrate limbs. Nature. 2002;416(6880):527–31. doi: 10.1038/416527a.

74. Jung H, Baek M, D’Elia KP, Boisvert C, Currie PD, Tay B-H, et al. The Ancient Origins of Neural Substrates for Land Walking. Cell. 2018;172(4):667–82. e15. doi: 10.1016/j.cell.2018.01.013.

75. Zdral S, Bordignon SG, Meyer A, Ros MA, Woltering JM. Dorsoventral limb patterning in paired appendages emerged via regulatory repurposing of an ancestral posterior fin module. Molecular Biology and Evolution. 2025;43(1). doi: 10.1093/molbev/msaf331.

76. McMahon C, Gestri G, Wilson SW, Link BA. Lmx1b is essential for survival of periocular mesenchymal cells and influences Fgf-mediated retinal patterning in zebrafish. Developmental Biology. 2009;332(2):287–98. doi: 10.1016/j.ydbio.2009.05.577.

77. O’Hara FP, Beck E, Barr LK, Wong LL, Kessler DS, Riddle RD. Zebrafish Lmx1b.1 and Lmx1b.2 are required for maintenance of the isthmic organizer. Development. 2005;132(14):3163–73. doi: 10.1242/dev.01898.

78. Ocaña OH, Coskun H, Minguillón C, Murawala P, Tanaka EM, Galcerán J, et al. A right-handed signalling pathway drives heart looping in vertebrates. Nature. 2017;549(7670):86–90. doi: 10.1038/nature23454.

79. Sánchez-Iranzo H, Galardi-Castilla M, Minguillón C, Sanz-Morejón A, González-Rosa JM, Felker A, et al. Tbx5a lineage tracing shows cardiomyocyte plasticity during zebrafish heart regeneration. Nature Communications. 2018;9(1):428. doi: 10.1038/s41467-017-02650-6.

80. Relaix F, Rocancourt D, Mansouri A, Buckingham M. A Pax3/Pax7-dependent population of skeletal muscle progenitor cells. Nature. 2005;435(7044):948–53. doi: 10.1038/nature03594.

81. Seger C, Hargrave M, Wang X, Chai RJ, Elworthy S, Ingham PW. Analysis of Pax7 expressing myogenic cells in zebrafish muscle development, injury, and models of disease. Developmental Dynamics. 2011;240(11):2440–51. doi: 10.1002/dvdy.22745.

82. Mercader N. Early steps of paired fin development in zebrafish compared with tetrapod limb development. Development, Growth & Differentiation. 2007;49(6):421–37. doi: 10.1111/j.1440-169X.2007.00942.x.

83. Abe G, Ide H, Tamura K. Function of FGF signaling in the developmental process of the median fin fold in zebrafish. Developmental biology. 2007;304(1):355–66. doi: 10.1016/j.ydbio.2006.12.040.

84. Bailon-Zambrano R, Keating MK, Sales EC, Nichols AR, Gustafson GE, Hopkins CA, et al. The sclerotome is the source of the dorsal and anal fin skeleton and its expansion is required for median fin development. Development. 2024;151(24):dev203025. doi: 10.1242/dev.203025.

85. Mabee PM, Crotwell PL, Bird NC, Burke AC. Evolution of median fin modules in the axial skeleton of fishes. Journal of Experimental Zoology. 2002;294(2):77–90. doi: 10.1002/jez.10076.

86. Miyamoto K, Kawakami K, Tamura K, Abe G. Developmental independence of median fins from the larval fin fold revises their evolutionary origin. Scientific Reports. 2022;12(1):7521. doi: 10.1038/s41598-022-11180-1.

87. Parichy DM, Elizondo MR, Mills MG, Gordon TN, Engeszer RE. Normal table of postembryonic zebrafish development: Staging by externally visible anatomy of the living fish. Developmental Dynamics. 2009;238(12):2975–3015. doi: 10.1002/dvdy.22113.

88. Bemis W, Grande L. Early development of the actinopterygian head. I. External development and staging of the paddlefish Polyodon spathula. Journal of Morphology. 1992;213(1):47–83.

89. Inoue JG, Miya M, Lam K, Tay B-H, Danks JA, Bell J, et al. Evolutionary Origin and Phylogeny of the Modern Holocephalans (Chondrichthyes: Chimaeriformes): A Mitogenomic Perspective. Molecular Biology and Evolution. 2010;27(11):2576–86. doi: 10.1093/molbev/msq147.

90. Didier DA, LeClair EE, Vanbuskirk DR. Embryonic staging and external features of development of the Chimaeroid fish, Callorhinchus milii (Holocephali, Callorhinchidae). Journal of Morphology. 1998;236(1):25–47.

91. Jerve A, Johanson Z, Ahlberg P, Boisvert C. Embryonic development of fin spines in Callorhinchus milii (Holocephali); implications for chondrichthyan fin spine evolution. Evolution & Development. 2014;16(6):339–53. doi: 10.1111/ede.12104.

92. Mosimann C, Kaufman CK, Li P, Pugach EK, Tamplin OJ, Zon LI. Ubiquitous transgene expression and Cre-based recombination driven by the ubiquitin promoter in zebrafish. Development. 2011;138(1):169–77. doi: 10.1242/dev.059345.

93. Bird NC, Mabee PM. Developmental morphology of the axial skeleton of the zebrafish, Danio rerio (Ostariophysi: Cyprinidae). Developmental Dynamics. 2003;228(3):337–57.

94. Grandel H, Schulte-Merker S. The development of the paired fins in the zebrafish (Danio rerio). Mechanisms of development. 1998;79(1-2):99–120.

95. Brown AM, Fisher S, Kathryn Iovine M. Osteoblast maturation occurs in overlapping proximal-distal compartments during fin regeneration in zebrafish. Developmental dynamics: an official publication of the American Association of Anatomists. 2009;238(11):2922–8.

96. Frazer KA, Pachter L, Poliakov A, Rubin EM, Dubchak I. VISTA: computational tools for comparative genomics. Nucleic Acids Research. 2004;32(suppl_2):W273–W9. doi: 10.1093/nar/gkh458.

97. Braasch I, Gehrke AR, Smith JJ, Kawasaki K, Manousaki T, Pasquier J, et al. The spotted gar genome illuminates vertebrate evolution and facilitates human-teleost comparisons. Nature Genetics. 2016;48(4):427–37. doi: 10.1038/ng.3526.

98. Keivany Y. Osteological Features of Eurypterygian Pelvic Girdles. Iranian Journal of Science and Technology, Transactions A: Science. 2017;41(4):989–1002. doi: 10.1007/s40995-017-0324-8.

99. Koch A-K, Moritz T. The pelvic girdle in extant gonorynchiformes (Teleostei: Otomorpha). Zoomorphology. 2024;143(1):141–50. doi: 10.1007/s00435-023-00628-1.

100. Stiassny ML, Moore JA. A review of the pelvic girdle of acanthomorph fishes, with comments on hypotheses of acanthomorph intrarelationships. Zoological Journal of the Linnean Society. 1992;104(3):209–42. doi: 10.1111/j.1096-3642.1992.tb00923.x.

101. Sendell-Price AT, Tulenko FJ, Pettersson M, Kang D, Montandon M, Winkler S, et al. Low mutation rate in epaulette sharks is consistent with a slow rate of evolution in sharks. Nature Communications. 2023;14(1):6628. doi: 10.1038/s41467-023-42238-x.

102. Feregrino C, Sacher F, Parnas O, Tschopp P. A single-cell transcriptomic atlas of the developing chicken limb. BMC Genomics. 2019;20(1):401. doi: 10.1186/s12864-019-5802-2.

103. He S, Wang L-H, Liu Y, Li Y-Q, Chen H-T, Xu J-H, et al. Single-cell transcriptome profiling of an adult human cell atlas of 15 major organs. Genome Biology. 2020;21(1):294. doi: 10.1186/s13059-020-02210-0.

104. Hirsinger E, Blavet C, Bonnin M-A, Bellenger L, Gharsalli T, Duprez D. Limb connective tissue is organized in a continuum of promiscuous fibroblast identities during development. iScience. 2024;27(7). doi: 10.1016/j.isci.2024.110305.

105. Kelly NH, Huynh NPT, Guilak F. Single cell RNA-sequencing reveals cellular heterogeneity and trajectories of lineage specification during murine embryonic limb development. Matrix Biology. 2020;89:1–10. doi: 10.1016/j.matbio.2019.12.004.

106. Stuart T, Srivastava A, Madad S, Lareau CA, Satija R. Single-cell chromatin state analysis with Signac. Nature Methods. 2021;18(11):1333–41. doi: 10.1038/s41592-021-01282-5.

107. Ma S, Zhang B, LaFave LM, Earl AS, Chiang Z, Hu Y, et al. Chromatin Potential Identified by Shared Single-Cell Profiling of RNA and Chromatin. Cell. 2020;183(4):1103–16.e20. doi: 10.1016/j.cell.2020.09.056.

108. Pliner HA, Packer JS, McFaline-Figueroa JL, Cusanovich DA, Daza RM, Aghamirzaie D, et al. Cicero Predicts *cis*-Regulatory DNA Interactions from Single-Cell Chromatin Accessibility Data. Molecular Cell. 2018;71(5):858–71.e8. doi: 10.1016/j.molcel.2018.06.044.

109. Rouco R, Rauseo A, Darbellay F, Sapin G, Bompadre O, Lopez-Delisle L, et al. Temporal constraints on enhancer usage shape the regulation of limb gene transcription. Nature Communications. 2026;17(1):5. doi: 10.1038/s41467-025-66055-6.

110. Gu WXW, Kania A. Identification of genes controlled by LMX1B in E13.5 mouse limbs. Developmental Dynamics. 2010;239(8):2246–55. doi: 10.1002/dvdy.22357.

111. Neufeld SJ, Wang F, Cobb J. Genetic Interactions Between Shox2 and Hox Genes During the Regional Growth and Development of the Mouse Limb. Genetics. 2014;198(3):1117–26. doi: 10.1534/genetics.114.167460.

112. Ye W, Song Y, Huang Z, Osterwalder M, Ljubojevic A, Xu J, et al. A unique stylopod patterning mechanism by Shox2-controlled osteogenesis. Development. 2016;143(14):2548–60. doi: 10.1242/dev.138750.

113. Yu L, Liu H, Yan M, Yang J, Long F, Muneoka K, et al. Shox2 is required for chondrocyte proliferation and maturation in proximal limb skeleton. Developmental Biology. 2007;306(2):549–59. doi: 10.1016/j.ydbio.2007.03.518.

114. Cobb J, Dierich A, Huss-Garcia Y, Duboule D. A mouse model for human short-stature syndromes identifies *Shox2* as an upstream regulator of *Runx2* during long-bone development. Proceedings of the National Academy of Sciences. 2006;103(12):4511–5. doi: 10.1073/pnas.0510544103.

115. Singh PNP, Ray A, Azad K, Bandyopadhyay A. A comprehensive mRNA expression analysis of developing chicken articular cartilage. Gene Expression Patterns. 2016;20(1):22–31. doi: 10.1016/j.gep.2015.11.001.

116. Singh PNP, Yadav US, Azad K, Goswami P, Kinare V, Bandyopadhyay A. NFIA and GATA3 are crucial regulators of embryonic articular cartilage differentiation. Development. 2018;145(2). doi: 10.1242/dev.156554.

117. Gao Y, Lan Y, Liu H, Jiang R. The zinc finger transcription factors Osr1 and Osr2 control synovial joint formation. Developmental Biology. 2011;352(1):83–91. doi: 10.1016/j.ydbio.2011.01.018.

118. Stricker S, Brieske N, Haupt J, Mundlos S. Comparative expression pattern of Odd-skipped related genes Osr1 and Osr2 in chick embryonic development. Gene Expression Patterns. 2006;6(8):826–34. doi: 10.1016/j.modgep.2006.02.003.

119. Suemoto H, Muragaki Y, Nishioka K, Sato M, Ooshima A, Itoh S, et al. Trps1 regulates proliferation and apoptosis of chondrocytes through Stat3 signaling. Developmental Biology. 2007;312(2):572–81. doi: 10.1016/j.ydbio.2007.10.001.

120. He D, Zhang M, Li Y, Liu F, Ban B. Insights into the ANKRD11 variants and short-stature phenotype through literature review and ClinVar database search. Orphanet Journal of Rare Diseases. 2024;19(1):292. doi: 10.1186/s13023-024-03301-y.

121. Wang S, Liao Y, Zhang H, Jiang Y, Peng Z, Ren R, et al. Tcf12 is required to sustain myogenic genes synergism with MyoD by remodelling the chromatin landscape. Communications Biology. 2022;5(1):1201. doi: 10.1038/s42003-022-04176-0.

122. Badia-i-Mompel P, Wessels L, Müller-Dott S, Trimbour R, Ramirez Flores RO, Argelaguet R, et al. Gene regulatory network inference in the era of single-cell multi-omics. Nature Reviews Genetics. 2023;24(11):739–54. doi: 10.1038/s41576-023-00618-5.

123. Kim D, Tran A, Kim HJ, Lin Y, Yang JYH, Yang P. Gene regulatory network reconstruction: harnessing the power of single-cell multi-omic data. npj Systems Biology and Applications. 2023;9(1):51. doi: 10.1038/s41540-023-00312-6.

124. Fleck JS, Jansen SMJ, Wollny D, Zenk F, Seimiya M, Jain A, et al. Inferring and perturbing cell fate regulomes in human brain organoids. Nature. 2023;621(7978):365–72. doi: 10.1038/s41586-022-05279-8.

125. Qiu Q, Chen H, Johnson RL. Lmx1b-expressing cells in the mouse limb bud define a dorsal mesenchymal lineage compartment. genesis. 2009;47(4):224–33. doi: 10.1002/dvg.20430.

126. Nakamura T, Gehrke AR, Lemberg J, Szymaszek J, Shubin NH. Digits and fin rays share common developmental histories. Nature. 2016;537(7619):225–8. doi: 10.1038/nature19322.

127. Tulenko FJ, Massey JL, Holmquist E, Kigundu G, Thomas S, Smith SM, et al. Fin-fold development in paddlefish and catshark and implications for the evolution of the autopod. Proceedings of the Royal Society B: Biological Sciences. 2017;284(1855):20162780. doi: 10.1098/rspb.2016.2780.

128. Freitas R, Gómez-Marín C, Wilson Jonathan M, Casares F, Gómez-Skarmeta José L. *Hoxd13* Contribution to the Evolution of Vertebrate Appendages. Developmental Cell. 2012;23(6):1219–29. doi: 10.1016/j.devcel.2012.10.015.

129. Yano T, Abe G, Yokoyama H, Kawakami K, Tamura K. Mechanism of pectoral fin outgrowth in zebrafish development. Development. 2012;139(16):2916–25. doi: 10.1242/dev.075572.

130. Zhang J, Wagh P, Guay D, Sanchez-Pulido L, Padhi BK, Korzh V, et al. Loss of fish actinotrichia proteins and the fin-to-limb transition. Nature. 2010;466(7303):234–7. doi: 10.1038/nature09137.

131. Lalonde RL, Moses D, Zhang J, Cornell N, Ekker M, Akimenko MA. Differential actinodin1 regulation in zebrafish and mouse appendages. Developmental Biology. 2016;417(1):91–103. doi: 10.1016/j.ydbio.2016.05.019.

132. Masselink W, Cole NJ, Fenyes F, Berger S, Sonntag C, Wood A, et al. A somitic contribution to the apical ectodermal ridge is essential for fin formation. Nature. 2016;535(7613):542–6. doi: 10.1038/nature18953.

133. Tulenko FJ, Augustus GJ, Massey JL, Sims SE, Mazan S, Davis MC. HoxD expression in the fin-fold compartment of basal gnathostomes and implications for paired appendage evolution. Scientific Reports. 2016;6(1):22720. doi: 10.1038/srep22720.

134. Hawkins MB, Zdral S, Naranjo S, Juliá M, Sánchez-Martín M, Daane JM, et al. Establishment, conservation, and innovation of dorsal determination mechanisms during the evolution of vertebrate paired appendages. bioRxiv. 2025:2025.07.02.662459. doi: 10.1101/2025.07.02.662459.

135. Logan C, Hornbruch A, Campbell I, Lumsden A. The role of Engrailed in establishing the dorsoventral axis of the chick limb. Development. 1997;124(12):2317–24. doi: 10.1242/dev.124.12.2317.

136. Allou L, Balzano S, Magg A, Quinodoz M, Royer-Bertrand B, Schöpflin R, et al. Non-coding deletions identify Maenli lncRNA as a limb-specific En1 regulator. Nature. 2021;592(7852):93–8. doi: 10.1038/s41586-021-03208-9.

137. Wurst W, Auerbach AB, Joyner AL. Multiple developmental defects in Engrailed-1 mutant mice: an early mid-hindbrain deletion and patterning defects in forelimbs and sternum. Development. 1994;120(7):2065–75. doi: 10.1242/dev.120.7.2065.

138. Monteiro A. Gene regulatory networks reused to build novel traits. BioEssays. 2012;34(3):181–6. doi: 10.1002/bies.201100160.

139. Ermakova GV, Meyntser IV, Lyubetsky VA, Zaraisky AG, Bayramov AV. The subfunctionalization of shox and shox2 paralogs in shark highlights both shared and distinct developmental mechanisms of branchial arches and fins. Frontiers in Cell and Developmental Biology. 2025;Volume 13 - 2025. doi: 10.3389/fcell.2025.1667637.

140. Freitas R, Zhang G, Cohn MJ. Biphasic Hoxd Gene Expression in Shark Paired Fins Reveals an Ancient Origin of the Distal Limb Domain. PLOS ONE. 2007;2(8):e754. doi: 10.1371/journal.pone.0000754.

141. Marlétaz F, de la Calle-Mustienes E, Acemel RD, Paliou C, Naranjo S, Martínez-García PM, et al. The little skate genome and the evolutionary emergence of wing-like fins. Nature. 2023;616(7957):495–503. doi: 10.1038/s41586-023-05868-1.

142. Nakamura T, Klomp J, Pieretti J, Schneider I, Gehrke AR, Shubin NH. Molecular mechanisms underlying the exceptional adaptations of batoid fins. Proceedings of the National Academy of Sciences. 2015;112(52):15940–5. doi: 10.1073/pnas.1521818112.

143. Onimaru K, Kuraku S, Takagi W, Hyodo S, Sharpe J, Tanaka M. A shift in anterior–posterior positional information underlies the fin-to-limb evolution. eLife. 2015;4:e07048. doi: 10.7554/eLife.07048.

144. Onimaru K, Marcon L, Musy M, Tanaka M, Sharpe J. The fin-to-limb transition as the re-organization of a Turing pattern. Nature Communications. 2016;7(1):11582. doi: 10.1038/ncomms11582.

145. Onimaru K, Tatsumi K, Tanegashima C, Kadota M, Nishimura O, Kuraku S. Developmental hourglass and heterochronic shifts in fin and limb development. eLife. 2021;10:e62865. doi: 10.7554/eLife.62865.

146. Barry SN, Crow KD. The role of HoxA11 and HoxA13 in the evolution of novel fin morphologies in a representative batoid (Leucoraja erinacea). EvoDevo. 2017;8(1):24. doi: 10.1186/s13227-017-0088-4.

147. O’Shaughnessy KL, Dahn RD, Cohn MJ. Molecular development of chondrichthyan claspers and the evolution of copulatory organs. Nature Communications. 2015;6(1):6698. doi: 10.1038/ncomms7698.

148. Matsuura M, Nishihara H, Onimaru K, Kokubo N, Kuraku S, Kusakabe R, et al. Identification of four Engrailed genes in the Japanese lamprey, Lethenteron japonicum. Developmental Dynamics. 2008;237(6):1581–9. doi: 10.1002/dvdy.21552.

149. McQueen E, Rebeiz M. Chapter Twelve - On the specificity of gene regulatory networks: How does network co-option affect subsequent evolution? In: Peter IS, editor. Current Topics in Developmental Biology. 139: Academic Press; 2020. p. 375–405.

150. Burke AC, Nowicki JL. A New View of Patterning Domains in the Vertebrate Mesoderm. Developmental Cell. 2003;4(2):159–65. doi: 10.1016/S1534-5807(03)00033-9.

151. Johanson Z. Evolution of paired fins and the lateral somitic frontier. Journal of Experimental Zoology Part B: Molecular and Developmental Evolution. 2010;314B(5):347–52. doi: 10.1002/jez.b.21343.

152. Shearman RM, Burke AC. The lateral somitic frontier in ontogeny and phylogeny. Journal of Experimental Zoology Part B: Molecular and Developmental Evolution. 2009;312B(6):603–12. doi: 10.1002/jez.b.21246.

153. Oliver JC, Tong X-L, Gall LF, Piel WH, Monteiro A. A Single Origin for Nymphalid Butterfly Eyespots Followed by Widespread Loss of Associated Gene Expression. PLOS Genetics. 2012;8(8):e1002893. doi: 10.1371/journal.pgen.1002893.

154. Westerfield M. The Zebrafish Book; A guide for the laboratory use of zebrafish (Danio rerio). 4th ed: University of Oregon Press; 2000.

155. Boisvert CA, Martins CL, Edmunds AG, Cocks J, Currie P. Capture, transport, and husbandry of elephant sharks (Callorhinchus milii) adults, eggs, and hatchlings for research and display. Zoo Biology. 2015;34(1):94–8. doi: 10.1002/zoo.21183.

156. Ballard W, Needham R. Normal embryonic stages of Polyodon spathula (Walbaum). Journal of Morphology. 1964;114(3):465–77.

157. Ballard WW, Mellinger J, Lechenault H. A series of normal stages for development of Scyliorhinus canicula, the lesser spotted dogfish (Chondrichthyes: Scyliorhinidae). Journal of Experimental Zoology. 1993;267(3):318–36.

158. Labun K, Montague TG, Krause M, Torres Cleuren YN, Tjeldnes H, Valen E. CHOPCHOP v3: expanding the CRISPR web toolbox beyond genome editing. Nucleic Acids Research. 2019;47(W1):W171–W4. doi: 10.1093/nar/gkz365.

159. Gagnon JA, Valen E, Thyme SB, Huang P, Ahkmetova L, Pauli A, et al. Efficient Mutagenesis by Cas9 Protein-Mediated Oligonucleotide Insertion and Large-Scale Assessment of Single-Guide RNAs. PLOS ONE. 2014;9(5):e98186. doi: 10.1371/journal.pone.0098186.

160. Modrell MS, Bemis WE, Northcutt RG, Davis MC, Baker CV. Electrosensory ampullary organs are derived from lateral line placodes in bony fishes. Nature Communications. 2011;2(1):496. doi: 10.1038/ncomms1502.

161. Suppermpool A, Trivedi C, Powell GT, Andrés M. Protocol for multiplex whole-mount RNA fluorescence *in situ* hybridization combined with immunohistochemistry in the mosquito brain. STAR Protocols. 2025;6(4). doi: 10.1016/j.xpro.2025.104109.

